# Location, location, location– choice of Voxel-Based Morphometry processing pipeline drives variability in the location of neuroanatomical brain markers

**DOI:** 10.1101/2021.03.09.434531

**Authors:** Xinqi Zhou, Renjing Wu, Yixu Zeng, Ziyu Qi, Stefania Ferraro, Shuxia Yao, Keith M. Kendrick, Benjamin Becker

## Abstract

Fundamental and clinical neuroscience has benefited from the development of automated computational analyses of Magnetic Resonance Imaging (MRI) data, such as Voxel-based Morphometry (VBM). VBM determines regional gray matter variations with high spatial resolution and results are commonly interpreted in a regional-specific manner, for instance with respect to which specific brain regions differ in volume between women and men. In excess of 600 papers using VBM are now published every year and a number of different automated VBM processing pipelines are frequently used in analyses although it remains to be fully and systematically assessed whether they come up with the same answers. Here we have therefore examined variability between four commonly used VBM pipelines in two large brain structural datasets. Spatial similarity, reproducibility and reliability of the processed gray matter brain maps was generally low between pipelines. Examination of sex-differences and age-related changes in gray matter volumes revealed considerable differences between the pipelines in terms of the specific regions identified as well as meta-analytic characterization of their function. In contrast, applying machine learning-based multivariate analyses allowed an accurate prediction of sex or age based on the gray matter maps across pipelines, although prediction accuracy differed strongly between them. Together the findings suggest that the choice of pipeline alone leads to considerable variability in brain structural analyses which poses a serious challenge for reproducibility as well as interpretation.

## Introduction

Human fundamental and clinical neuroscience aims to determine the contribution of specific brain systems to mental processes and to identify which specific systems exhibit pathological alterations in mental and neurological disorders. Neuroimaging techniques have become a widely employed tool to this end and due to their high spatial resolution and non-invasive nature, Magnetic Resonance Imaging (MRI)-based assessments of brain structure and function have become one of the most widely used neuroimaging techniques. However, the complexity and flexibility of workflows in MRI analyses and differences between the handful of commonly used analysis software packages may lead to high variability in neuroimaging results (Poldrack et al., 2017), thus challenging the interpretation in terms of the precise mapping of mental processes as well as regional-specific biomarkers for brain disorders. Compared to the processing of functional imaging data (fMRI), brain morphometry analyses of T1-weighted structural images may allow less processing variations and may have higher test-retest reliability (Buimer et al., 2020; Madan & Kensinger, 2017; Melzer et al., 2020; Poldrack et al., 2017; Shou et al., 2013), however, the choice of the analytic software may still have a considerable impact on results obtained in brain structural analyses. The variability in terms of whether and which specific brain regions pass the statistical threshold in turn strongly impacts the interpretation of findings with respect to structure-function mapping or brain-based biomarkers and could significantly impedes the sensitivity of subsequent neuroimaging meta-analyses which aim to determine robust brain structural markers across a number of original studies.

Neuroanatomical research has tremendously benefited from the development of automated computational approaches such as Voxel-based Morphometry (VBM), probing differences in regional grey matter volumes between groups, and the more recent surface/volume-based approaches, investigating grey matter absolute metrics (e.g. cortical thickness and subcortical volumes). VBM represents the most widely used brain structural analytic approach to date (e.g. a simple literature search using the term “voxel-based morphometry” or “VBM” on PubMed revealed 6,210 studies, https://pubmed.ncbi.nlm.nih.gov, from 1993 to Nov 19^th^, 2020). The standardized and highly automated VBM workflow includes segmentation of gray matter from other brain tissues, normalization into standard stereotactic space and smoothing with a Gaussian kernel before inferential statistics being applied. The corresponding inferential voxel-wise statistical models most commonly determine (1) differences in regional gray matter volume (GMV) between groups, e.g. by comparing patients and controls or men and women (e.g. Becker et al., 2015; Lotze et al., 2019; Ritchie et al., 2018; Ruigrok et al., 2014), or (2) associations between variations in regional GMV and behavioral phenotypes, including individual variations in learning, age, or disorder-relevant traits (Llera et al., 2019; Miller et al., 2016; Nostro et al., 2017; Taubert et al., 2011; Zhao et al., 2021; Zhou et al., 2020). Significant associations and differences are commonly interpreted in a regional-specific fashion, e.g. mapping specific behavioral functions to specific brain systems, determining which brain regions undergo age-related changes or which regions contribute to mental disorders. More recently, machine learning-based multivariate analytic approaches such as Multivariate Pattern Analyses (MVPA) have been increasingly applied to VBM data to detect subtle and spatially distributed patterns of brain structural variations, particularly to improve biomarker-based diagnostics of mental disorders (Nunes et al., 2020; Premi et al., 2016; Wang et al., 2016).

A number of software packages have been developed and are widely applied for VBM analyses. Among them the currently most widely used ones are the Computation Anatomy Toolbox (CAT, www.neuro.uni-jena.de/cat), which is implemented in Statistical Parametric Mapping (SPM, https://www.fil.ion.ucl.ac.uk/spm/software/spm12/), and FSLVBM and FSLANAT, which are based on the FMRIB Software Library (FSL, https://fsl.fmrib.ox.ac.uk). Recently, to enhance the robustness and reproducibility of neuroimaging analyses, new automated preprocessing pipelines for structural MRI (e.g., sMRIPrep, https://www.nipreps.org/smriprep/) have been developed. Although the software packages generally employ similar processing steps to volumetric T1-weighted (anatomical) MRI data, differences in specific processing steps and their implementation exist, raising the question of whether the choice of the specific software and the application of the software-specific default processing configurations may lead to variability in the results obtained, which makes it crucial to robustly test the reliability and replicability of the employed tools.

A recent study examined reliability and replicability in cortical thickness measures using different software packages in large datasets of healthy subjects and reported a similar cortical thickness distribution across software packages, although the absolute estimated values varied considerably among pipelines (Kharabian Masouleh et al., 2020). In contrast, studies exploring the replicability of VBM in samples of neurological patients revealed considerable variations among the processing pipelines and results suggesting that the VBM processing pipeline strongly affects the clinical interpretation (Popescu et al., 2016; Rajagopalan & Pioro, 2015). Specifically, spatial normalization inaccuracies and different spatial normalization templates and methods challenge one of the main assumptions of VBM, namely that individual brain differences and anatomical correspondence of brain areas are maintained during the spatial normalization process (Bookstein, 2001; Mechelli et al., 2005; Pepe et al., 2014; Senjem et al., 2005; Shen et al., 2007). More importantly, VBM lacks a clear in-vivo or ex-vivo histological neurobiological validation in humans (Pepe et al., 2014). From a more general perspective, these observations strongly challenge the interpretation of VBM findings as plausible biomarkers for disorders or phenotypical variations.

Against this background the present study systematically examined the influence of the choice of the most commonly processing software packages on VBM results (FSLVBM and FSLANAT as implemented in FSL v6.0, sMRIPrep 0.6.2, and CAT 12.7, all recent releases). To model the common scientific workflow while controlling for pathology-related brain structural variations the recommended default configurations were employed to determine between group-differences and biological associations within two independent samples of healthy individuals (n = 200; n = 494). Given the previously reported low robustness of associations between latent variables and structural brain indices (see Kharabian Masouleh et al., 2019) our analyses focused on biological variables which have demonstrated greater robustness and thus potential biological validity (Fjell et al., 2014; Kharabian Masouleh et al., 2016; Lotze et al., 2019; Marek et al., 2020; Ritchie et al., 2018; Willette & Kapogiannis, 2015).

To determine the effects of the choice of processing pipeline on the results of a typical VBM study, we therefore examined sex-differences and age-related changes with univariate analyses (group-differences and regression, respectively) as well as multivariate analyses (machine learning based MVPA) in two comparably large datasets after processing with the commonly used VBM pipelines. Specifically, the following systematic analytic steps were conducted. First, spatial similarity and intraclass correlation (ICC, both voxel-wise and image-based estimations) were examined to determine the homogeneity and reliability of outcomes across pipelines before statistical analyses. Second, results with respect to sex differences in GMV from univariate between-groups comparisons between males and females were compared across pipelines. Third, results with respect to age-related GMV changes from univariate linear regression analysis were compared across pipelines. Finally, effects of pipeline on multivariate prediction accuracy were examined by MVPA-based predictions of sex and age based on whole-brain GMV maps across pipelines.

## Methods

### Datasets

Dataset 1 included T1-weigthed anatomical data from 200 healthy Chinese participants aged 18-26 years old (mean = 21.45 years old, SD = 2.18; 100 females and 100 males matched for age; sample details see also Liu et al. (2020)). This dataset served to determine variations across the four analytic pipelines with respect to determining gray matter differences in between-subject designs using the example of sex differences.

Dataset 2 included 494 healthy Chinese participants aged 19-80 years (mean = 45.18 years, SD = 17.44, 187 males) from an openly available dataset encompassing T1-weigthed anatomical and resting-state functional MRI data (details please see Wei et al. (2018)). This dataset served to determine variations between the software packages with respect to determining linear associations between biological indices and gray matter volume at the example of age-related changes. For detailed structural MRI acquisition parameters please see ***supplemental methods***.

### Data quality control

First, we inspected apparent artifacts and image quality by visual inspection which confirmed high image quality. Second, automated quality assessment by the MRIQC toolbox (https://mriqc.readthedocs.io/) (Esteban et al., 2017) was employed to further evaluate raw data quality, including signal-to-noise ratio (SNR), foreground to background energy ratio (FBER), percent of artefact voxels (Qi1) (details and results see ***supplemental material*** Fig. S1 and S2 and Wei et al. (2018)). Third, the CAT12.7 (r1720) (http://www.neuro.uni-jena.de/cat/) quality assurance (QA) framework for empirical quantification of quality differences across scans and studies was applied. This retrospective QA allows the evaluation of essential image parameters such as noise, inhomogeneities, and image resolution which can be integrated into a single quality index (dataset 1: mean = 81.68, SD = 1.61, range = 73.48-84.48; dataset 2: mean = 84.51, SD = 1.27, range = 76.9-86.24; scores >70 indicates satisfactory to excellent image quality).

### Preprocessing pipelines

VBM analyses commonly write out two types of structural indices, referred to as volume and concentration depending on whether a modulation step is employed or not (Ashburner & Friston, 2000; Gennatas et al., 2017; Good et al., 2001). In line with the advantages of and wider use of modulated images (volume) all subsequent analyses focused on modulated data.

Four separate preprocessing pipelines for the voxel-wise estimation of local grey matter volume were established in accordance with the default or recommended configurations in the respective manuals of the software packages (Fig. 1). One pipeline was based on CAT12.7 (r1720) (http://www.neuro.uni-jena.de/cat/) (CAT); two pipelines were based on FSL v6.0 (https://fsl.fmrib.ox.ac.uk/fsl/fslwiki/FSL, Jenkinson et al. (2012); Smith et al. (2004)) (FSLVBM and FSLANAT respectively); and one pipeline was based on sMRIPrep 0.6.2 (Esteban et al. (2019), RRID:SCR_016216, https://www.nipreps.org/smriprep/) (sMRIPrep).

**Fig. 1.**
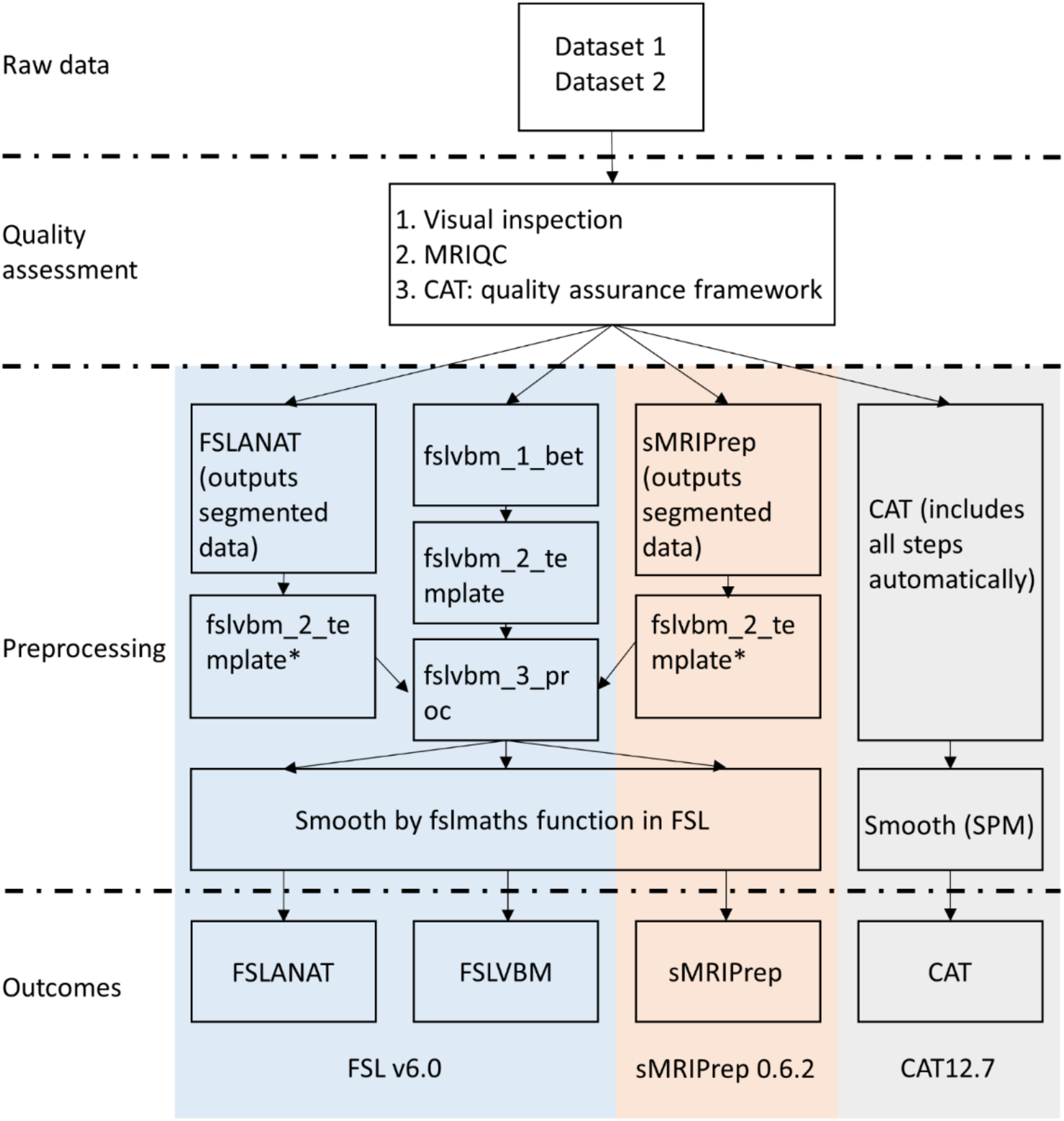
The overview flowchart of preprocessing steps across four pipelines. *Given that FSLANAT and sMRIPrep pipelines are mainly used for segmenting GM, WM, and CSF data, the segmented GM outcomes (in native space) were therefore subjected to preprocessing steps from fslvbm_2_template (except for segmentation) and fslvbm_3_proc to produce normalized and modulated GM data.

CAT12.7 was implemented in SPM12 v7219 (Welcome Department of Cognitive Neurology, London, UK, https://www.fil.ion.ucl.ac.uk/spm/software/spm12/). In general, the standard VBM preprocessing protocols of CAT12 (outlined in the CAT12.7 manual) were employed. Briefly, the T1-weighted images were bias-corrected, segmented into grey matter (GM), white matter (WM), and cerebrospinal fluid (CSF) and spatially normalized to the standard Montreal Neurological Institute (MNI) space using the ICBM152 template (East Asian, additional results obtained with the Caucasian template did not affect the results, see supplements Fig. S8) with voxel size 2×2×2 mm. GM images were smoothed with three Gaussian kernels with commonly used smoothing kernels (8 mm, 10 mm, and 12mm) at full-width at half maximum (FWHM) for subsequent statistical analysis and total intracranial volume (TIV) was estimated to correct for individual differences in brain size. Default parameters were applied unless indicated otherwise.

Two different default preprocessing pipelines were established in FSL (Jenkinson et al., 2012; Smith et al., 2004): (1) FSLANAT (https://fsl.fmrib.ox.ac.uk/fsl/fslwiki/fsl_anat), and (2) FSLVBM (https://fsl.fmrib.ox.ac.uk/fsl/fslwiki/FSLVBM). FSLANAT is a general pipeline for processing anatomical images encompassing the following steps. First, reorientation of all T1-weighted images to the standard MNI orientation and automatic cropping of the image. Second, bias-field correction for RF/B1-inhomogeneity-correction. Third, non-linear registration to standard space using FNIRT followed by brainextraction and tissue-type segmentation. Finally, a study-specific template is created (similar as in FSLVBM) and the GM images are registered to this template. The FSLVBM default pipeline included the following four steps: First, removal of nonbrain tissue using BET (fslvbm_1_bet). Second (fslvbm_2_template), tissue-type segmentation using the Automated Segmentation Tool (FAST), to segment the images into GM, WM, and CSF. Third, non-linear registration to the GM ICBM-152 template using the affine registration tool FNIRT, and next creation of a study-specific template. Finally, the GM images were non-linearly registered to the study-specific template using FNIRT (fslvbm_3_proc). The calculation of TIV for both FSLANAT and FSLVBM adhered to the protocols provided by the ENIGMA project (http://enigma.ini.usc.edu/protocols/imaging-protocols/protocol-for-brain-and-intracranial-volumes/#fsl).

sMRIPrep 0.6.2 (Esteban et al. (2019), RRID:SCR_016216, https://www.nipreps.org/smriprep/) is a structural MRI data preprocessing pipeline designed to provide an easily accessible, state-of-the-art interface that is robust to variations in scan acquisition protocols and that requires minimal user input, while providing easily interpretable and comprehensive error and output reporting. The workflow is based on Nipype 1.5.0 (Gorgolewski et al. (2011), RRID:SCR_002502). The similar workflow is also used in fMRIPrep anatomical preprocessing workflow (Esteban et al. (2019), https://fmriprep.org/). In the present study, the T1-weighted (T1w) image was corrected for intensity non-uniformity (INU) with N4BiasFieldCorrection (Tustison et al., 2010), distributed with ANTs 2.2.0 (Avants et al. (2008), RRID:SCR_004757), and used as T1w-reference throughout the workflow. The T1w-reference was then skull-stripped with a Nipype implementation of the antsBrainExtraction.sh workflow (from ANTs), using OASIS30ANTs as target template. Brain tissue segmentation of CSF, WM and GM was performed on the brain-extracted T1w using fast (FSL 5.0.9, RRID:SCR_002823, Zhang et al. (2001)). Considering no TIV estimation is provided by sMRIPrep, the corresponding brain size for the analysis was computed by summarizing the tissue types (GM+WM+CSF).

Given that FSLANAT and sMRIPrep pipelines are mainly used for segmenting GM, WM, and CSF data, the initial brain-extraction (fslvbm_1_bet) and segmentation (first part of fslvbm_2_template) stages were not included. In line with the recommendations in the user manuals the segmented GM outcomes (in native space) were therefore subjected to preprocessing steps from fslvbm_2_template and fslvbm_3_proc to produce modulated GM data.

To keep preprocessing consistent within each platform the fslmaths function was used to smooth FSL processed data (FSLVBM, FSLANAT and sMRIPrep) with comparable smoothing kernels (sigma = 3.5, 4.3, 5.2, approximately corresponding to FWHM - 3.5×2.3=8.05≈8, 4.3×2.3=9.89≈10, and 5.2×2.3=11.96≈12) as the CAT data. For the CAT preprocessing, SPM smoothing was conducted with FWHM = 8, 10, and 12 respectively.

## Analytic strategy

### Spatial similarity

Pearson’s correlation coefficients were employed to compute the spatial similarity of the modulated GM maps of the preprocessed data from the four pipelines for dataset 1 (male and female) and dataset 2, and at different smoothing kernels (Fig. 2).

**Fig. 2.**
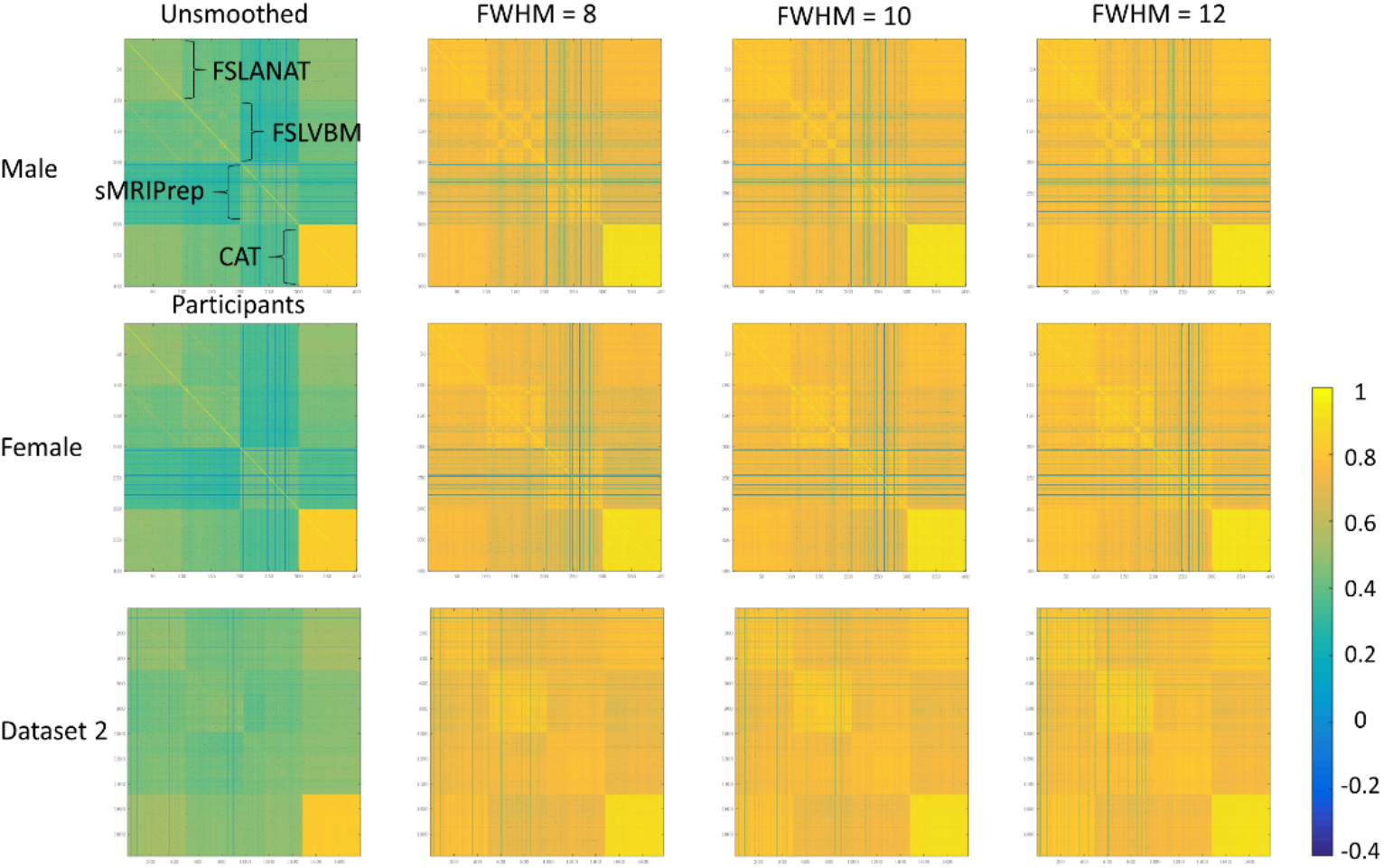
Spatial similar maps across the processing pipelines. Top and middle rows correspond to male and female samples separately from dataset 1. The bottom row shows data from the entire sample of dataset 2. Each column corresponds to a smoothing level (unsmoothed, 8, 10, and 12mm FWHM). The color grading reflect r values ranging from −0.4 to 1 (no r value was lower than −0.4). Each line of both x and y axes in each matric map refers to one participant.

Considering highly similar patterns across unsmoothed and the three smoothing kernels (FWHM 8, 10, and 12) further statistical analyses were only conducted with the z-transformed r values of the FWHM 8 smoothed data. Spatial similarities across the four pipelines were examined with respect to three indices: (1) between software and between participants, (2) between software and within participants, and (3) within software and between participants. For each comparison, ANOVAs were employed separately in both dataset 1 (male, female sample) and dataset 2 (Fig. 3, unsmoothed data please see ***supplemental material*** Fig. S3).

**Fig. 3.**
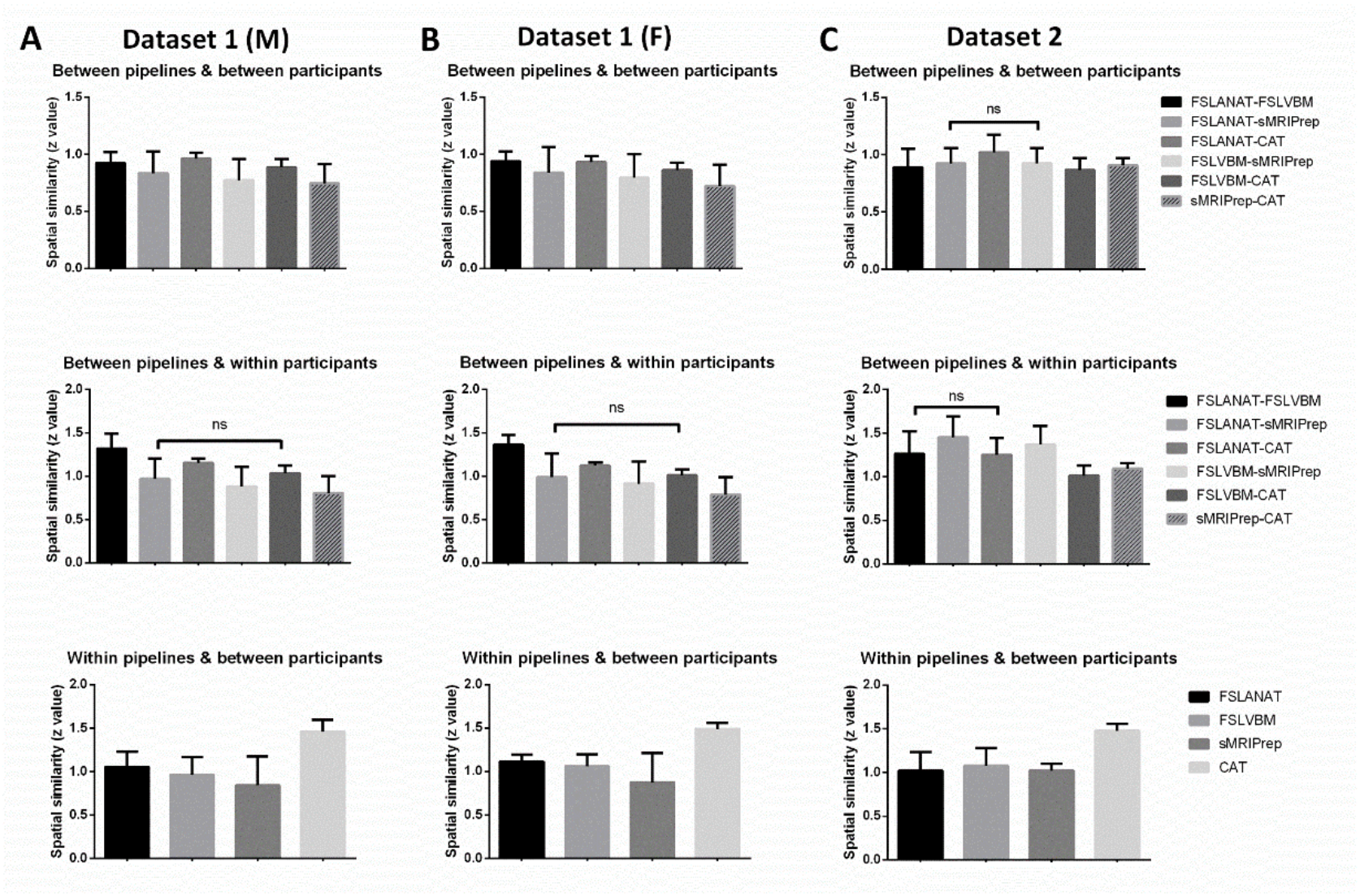
Mean similarity and standard deviation (SD) of different pipelines and pipeline pairs. Column A and B display the data from dataset 1 for males (A) and females (B) respectively. Column C displays the data from dataset 2. Post hoc tests were controlled for multiple comparison using a Bonferroni corrected p < 0.01. M = males, F = females.

### Reliability

Reliability across pipelines was evaluated using two approaches: (1) voxel-level univariate reliability was examined using the intraclass correlation coefficient (ICC) implemented by a linear mixed model in DPABI (Yan et al., 2016), and (2) wholebrain multivariate reliability, using the image intraclass correlation coefficient (I2C2), which represents a multivariate image measurement error model (Shou et al., 2013). ICC (ICC (3,1) with linear mixed models as used in the current study) and I2C2 can estimate the consistency between the different pipelines on the voxel and whole brain level respectively (Noble et al., 2019; Shou et al., 2013). The reliability between the datasets as assed by the coefficient is commonly interpreted as follows: <0.4 poor; 0.4 – 0.59 fair; 0.60 – 0.74 good; > 0.74 excellent (Cicchetti & Sparrow, 1981; Noble et al., 2019; Noble et al., 2017).

### Statistical analyses

#### Univariate analyses

To account for potential interactions between preprocessing and inferential statistical procedures all univariate analyses in the current study were conducted in SPM12 and across different multiple comparisons corrections, including classical statistical parameter tests (threshold at voxel-level *p* < 0.001, and cluster-level *p_FWE_* < 0.05 with initial cluster forming voxel-level *p* < 0.001 respectively) as well as threshold-free cluster enhancement (TFCE with 5,000 permutations, threshold at *p* < 0.001, and *p_FWE_* < 0.05 respectively). For completeness, the uncorrected voxel-level results (*p* < 0.001 and TFCE *p* < 0.001) are provided in ***supplemental material*** Fig. S4 and S5.

#### Between-group difference approach: sex differences univariate analyses

Independent sample t-tests were employed to determine significant differences in regional gray matter volume between men and women. Age and TIV were included in the models as recommended for VBM analyses to control for age- and global brain size related variations.

#### Association approach: age-related changes

Multiple linear regression models were employed to explore associations between age and regional GMV including sex and TIV as covariates.

#### Multivariate pattern analysis approach: prediction of sex and age

The state of art machine-learning framework in neuroimaging (Kohoutova et al., 2020) was adopted to explore whether use of the different pipelines will affect prediction accuracy of sex and age by means of distributed brain structural variations. For the categorical prediction (sex) the 200 healthy participants from dataset 1 were divided into two sex- and age-matched independent samples which served as training and test datasets respectively. A sex differences classifier based on a support vector machine (SVM) was trained on the training data with a bootstrapping test (5,000 permutations, *p_FDR_* < 0.05), evaluated by means of leave-one-out cross-validation (LOOCV) and subsequently tested on the independent sample to predict sex within- and between-pipelines. To estimate the effect size of each classification Cohen’s d for between-subjects designs was employed (Lakens, 2013). For prediction of a continuous variable (age) support vector regression (SVR) model with a bootstrapping test (5,000 permutations, *p_FDR_* < 0.05) and 5-fold crossvalidation was applied to dataset 2. Performance was next quantified by evaluation of correlation strengths between predicted and true age.

Of note, the aim of the MVPA was not to determine an optimized algorithm or feature set to predict sex or age but rather to determine whether different processing pipelines affect prediction accuracy and whether the GMV maps generally encode biologically meaningful information.

#### Data availability

Unthreshold statistical maps and pattern weight images are available on OSF (https://osf.io/p5b6f/). Other data can be obtained from the corresponding authors upon reasonable request.

## Results

### Spatial similarity within- and between-pipelines

Examination of the spatial similarity between pair-wised permutations of the four processing pipelines, i.e. FSLANAT-FSLVBM, FSLANAT-sMRIPrep, FSLANAT-CAT, FSLVBM-sMRIPrep, FSLVBM-CAT, and sMRIPrep-CAT and the within pipeline similarities by means of ANOVA models revealed significant main effects of pipeline pair or pipeline, respectively. Bonferroni-corrected post hoc tests for dataset 1 (male data presented in column A, female data presented in column B in Fig 3) demonstrated that each pipeline pair was significantly different from other pairs with respect to similarity between pipelines and between participants (Bonferroni’s corrected *p* < 0.01; Fig. 3A & 3B upper panel). A similar pattern of results was observed for the between pipelines and within participants analysis, except that the difference between FSLANAT-sMRIPrep and FSLVBM-CAT did not reach significance (Fig. 3A & 3B middle panel).

A similar pattern of post hoc results was observed for dataset 2 such that each pipeline and pipeline pair significantly differed from the others (Fig. 3C), except that there were no significant differences between FSLANAT-sMRIPrep and FSLVBM-sMRIPrep in the between-pipelines and between-participants comparisons, while FSLANAT-FSLVBM and FSLANAT-CAT did not significantly differ for the between-pipelines and within-participants comparison. For detailed ANOVA and post hoc results please see ***supplemental material*** results and Table S1-S9. These findings indicate significant spatial dissimilarities between GMV maps from the same participants between pipelines as well as from the same pipeline between participants. Specifically, across the datasets the lowest spatial similarity was observed between CAT and FSLVBM, and CAT and sMRIPrep respectively. Notably CAT reached a considerably higher within-pipeline spatial similarity as compared to the other pipelines (Fig. 3A & 3B & 3C bottom), reflecting a higher homogeneity of the processed GMV maps between participants when the data was processed with CAT.

### Reliability across pipelines

Examination of ICC maps for each between pipeline comparison revealed generally lower regional consistency between the GMV maps computed by different pipelines (see Fig. 4 for dataset 1 and 2). Consistencies on the voxel-level showed some regional variations, with between pipeline reliability being consistently low in parietal and frontal regions. An exception was a comparably high reliability within FSLANAT vs. FSLVBM of dataset 1, and FSLANAT vs. sMRIPrep of dataset 2 (Fig. 4). Of note, across the two datasets different pipelines exhibited relatively high between pipeline reliability which may be explained by the fact that the data was acquired in different imaging centers and thus suggesting that acquisition and pipeline differences interact to influence variability (see Yamashita et al., 2019).

**Fig. 4.**
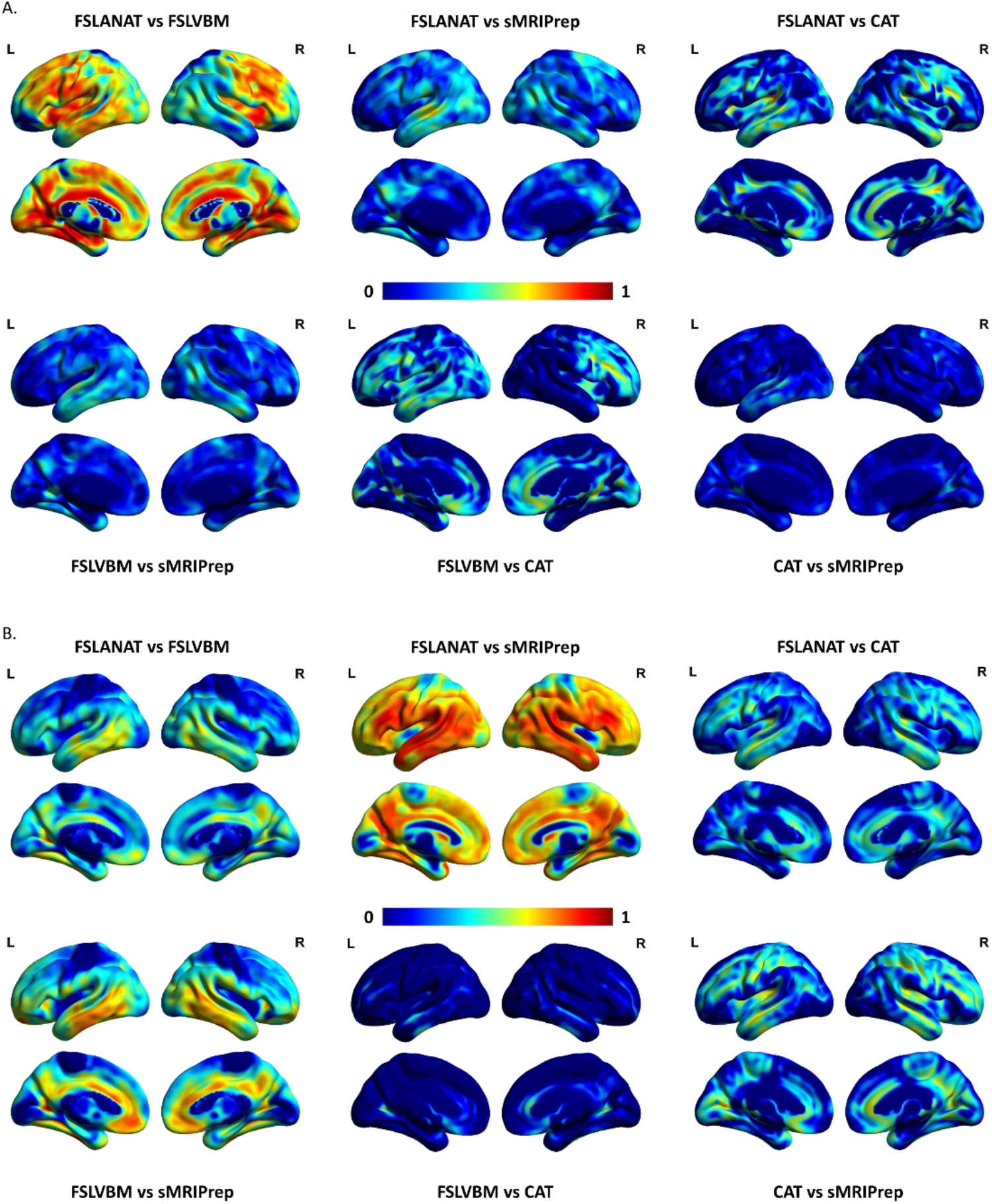
The voxel-level intraclass correlation coefficient (ICC) maps between pipelines of (A) dataset 1 and (B) dataset 2. L = left, R = right, and the color grading reflect the ICC value.

Examination of the I2C2 indices revealed a generally poor consistency between the pipelines (all image intraclass correlation coefficients < 0.4) suggesting low inter-pipeline reliability. Across the two datasets only reliability between CAT and FSLANAT or CAT and FSLVBM approached the ‘fair’ criterion (Table S11). For dataset 2, moderate higher I2C2 indices were observed between sMRIPrep and CAT, and across all pipelines compared to dataset 1.

### Between-group difference approach: sex differences univariate analyses

We mainly focused on the common and different brain areas of sex differences across the four pipelines. To this end, the percent of the common and different voxels in all significant voxels across the four pipelines was calculated (***supplemental methods***). For parametric statistics (cluster-level *p_FWE_* < 0.05) only 10.98% spatial overlap of the results for sex-differences among the three pipelines (FSLANAT, FSLVBM, and CAT, Table 1) were observed while the different pipelines mapped considerable pipeline-unique GMV sex-differences (up to 54.73% unique GMV sex differences identified by one, but not the other pipelines, Table 1). Between the pipelines overlap for male>female was observed in the lingual gyrus, precuneus, left hippocampus, bilateral parahippocampal cortex, olfactory cortex, left putamen, and left insula (Fig. 5A). While no common regions for female > male were observed among three of the pipelines, the FSL pipelines shared only 13.16% overlap (Table 1) with overlapping higher GMV for females being located in the bilateral postcentral cortex, right angular, right inferior parietal lobule, and cerebellum (Fig. 5A). In contrast to the few shared regions widespread variations in the location and extent of the identified GMV sex-differences were observed specifically in medial prefrontal and occipital regions. For instance, whereas CAT revealed higher GMV in widespread cerebellar and limbic regions in men, FSLANAT and FSLVBM revealed higher GMV in widespread posterior/superior parietal regions in women (Fig. 5A).

**Table 1.** Percent overlap of GMV sex-differences as revealed by the four pipelines

|  | Male>female |  | Female>male |  |
| --- | --- | --- | --- | --- |
|  | Parametric<sup>1</sup> | Non-parametric<sup>2</sup> | Parametric | Non-parametric |
| CAT (unique) | 54.73% | 38.94% | - | - |
| FSLVBM (unique) | 16.57% | 8.14% | 8.02% | 2.03% |
| FSLANAT (unique) | 1.38% | 0.29% | 78.82% | 88.64% |
| sMRIPrep (unique) | - | - | - | - |
| CAT ∩ FSLVBM | 3.76% | 9.84% | - | - |
| CAT ∩ FSLANAT | 4.55% | 10.93% | - | - |
| CAT ∩ sMRIPrep | - | - | - | - |
| FSLVBM ∩ FSLANAT | 8.02% | 12.97% | 13.16% | 9.33% |
| FSLVBM ∩ sMRIPrep | - | - | - | - |
| FSLANAT ∩ sMRIPrep | - | - | - | - |
| CAT ∩ FSLVBM ∩ FSLANAT | 10.98% | 18.89% | - | - |
| CAT ∩ FSLVBM ∩ sMRIPrep | - | - | - | - |
| CAT ∩ FSLANAT ∩ sMRIPrep | - | - | - | - |
| FSLVBM ∩ FSLANAT ∩ sMRIPrep | - | - | - | - |
| CAT ∩ FSLVBM ∩ FSLANAT ∩ sMRIPrep | - | - | - | - |

<sup>1</sup>cluster-level $p_{FWE} < 0.05$
<sup>2</sup>TFCE $p_{FWE} < 0.05$

**Fig. 5.**
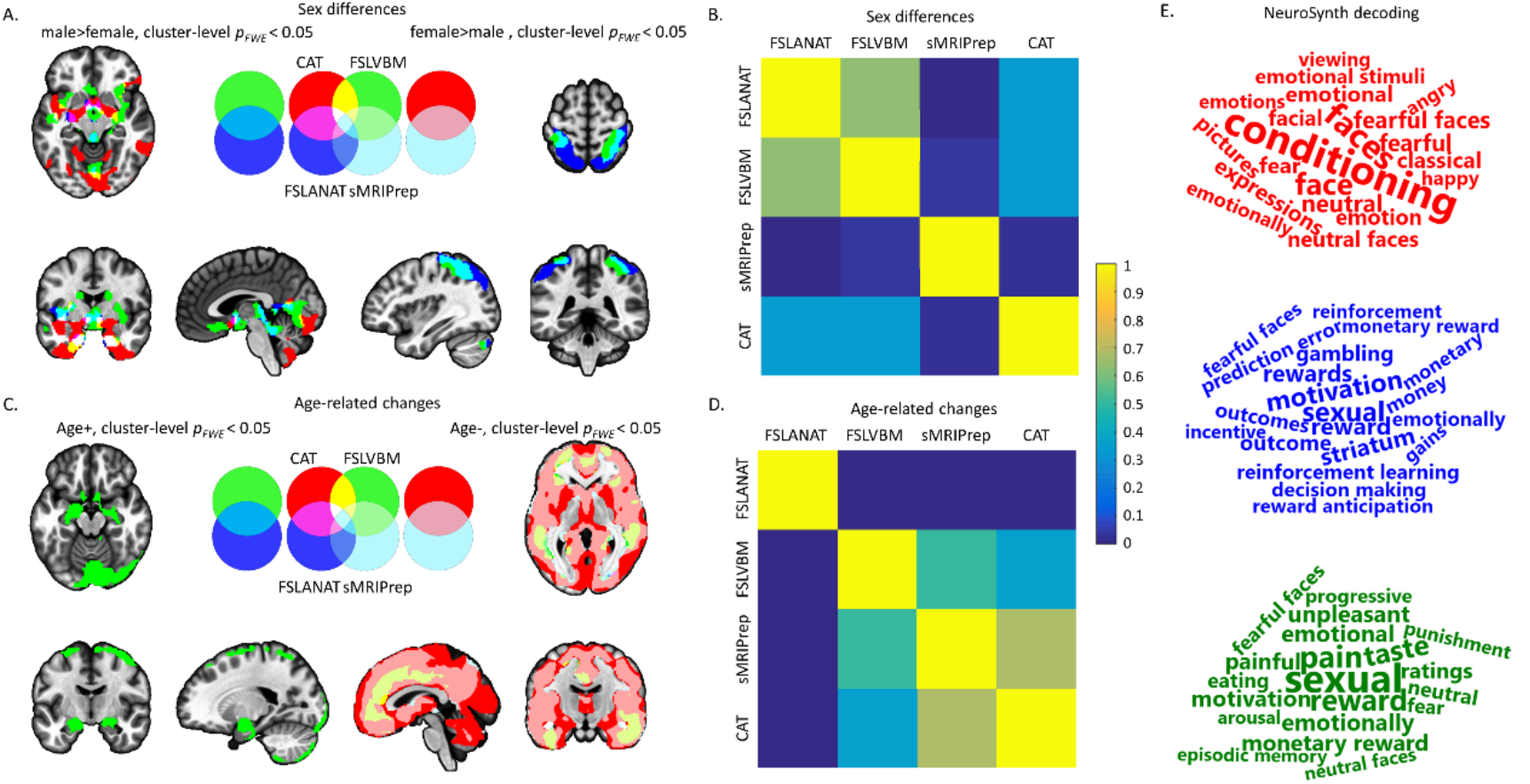
Similarities and dissimilarities between the pipelines with respect to determining GMV sex differences and age-related GMV changes. Fig. 5A and C present results from parametric statistic overlaps at cluster-level *p_FWE_* < 0.05 with initial cluster forming voxel-level *p* < 0.001. The left panels of A display results for the male>female contrast. The right panels of A correspond to the female>male contrast. The left panels of C depict brain regions with increasing GMV with age. The right panels of C depict decreases with age. For A and C the pipelines are coded as: red = CAT, green = FSLVBM, blue = FSLANAT, light blue = sMRIPrep, additional colors visualize the overlap between the results. B and D represent the variability of unthresholded statistical maps. The correlation values between wholebrain unthresholded statistical maps of four pipelines were computed respectively for (B) sex differences, and (D) age-related effects. Only positive values are visualized for display purpose. E, decoding the functional properties of the identified brain regions of male>female (A, red = CAT, green = FSLVBM, blue = FSLANAT) using NeuroSynth. Only the top 20 terms are visualized. The font size reflects the size of the correlation.

Results from the non-parametric statistics (TFCE *p_FWE_* < 0.05) were highly similar to the parametric statistic results, suggesting that the pipeline differences are robust across statistic models (details please see ***supplemental results***). Notably, in some instances the overlap between the software packages increased slightly using the non-parametric approach (Table 1 and Fig. S6A).

To further account for potential interaction effects between the preprocessing pipelines and correction for multiple comparisons, correlations between unthresholded statistical between-group difference maps across the four pipelines were computed (similar approach see Botvinik-Nezer et al., 2020). The spatial pattern of similarities of sex-dependent GMV variations ranged from −0.0033 to 0.6328 (Fig. 5B), with a particular low spatial overlap of sex-differences revealed by sMRIPrep with those obtained by other pipelines, and results revealed by CAT being very dissimilar from sex-differences obtained by the FSL pipelines. Finally, FSLANAT and FSLVBM had the highest similarity between the unthresholded maps.

### Prediction approach: sex differences in multivariate pattern analyses

Initially pipeline-specific classifiers were developed on the training dataset and benchmarked in the independent dataset preprocessed by the identical pipeline. In general, classifiers developed on each pipeline accurately predicted sex in the independent data (accuracy ranging from 68% (sMRIPrep) to 94% (CAT), Cohen’s d = 0.2967 to 2.2815, Fig. 6B).

**Fig. 6.**
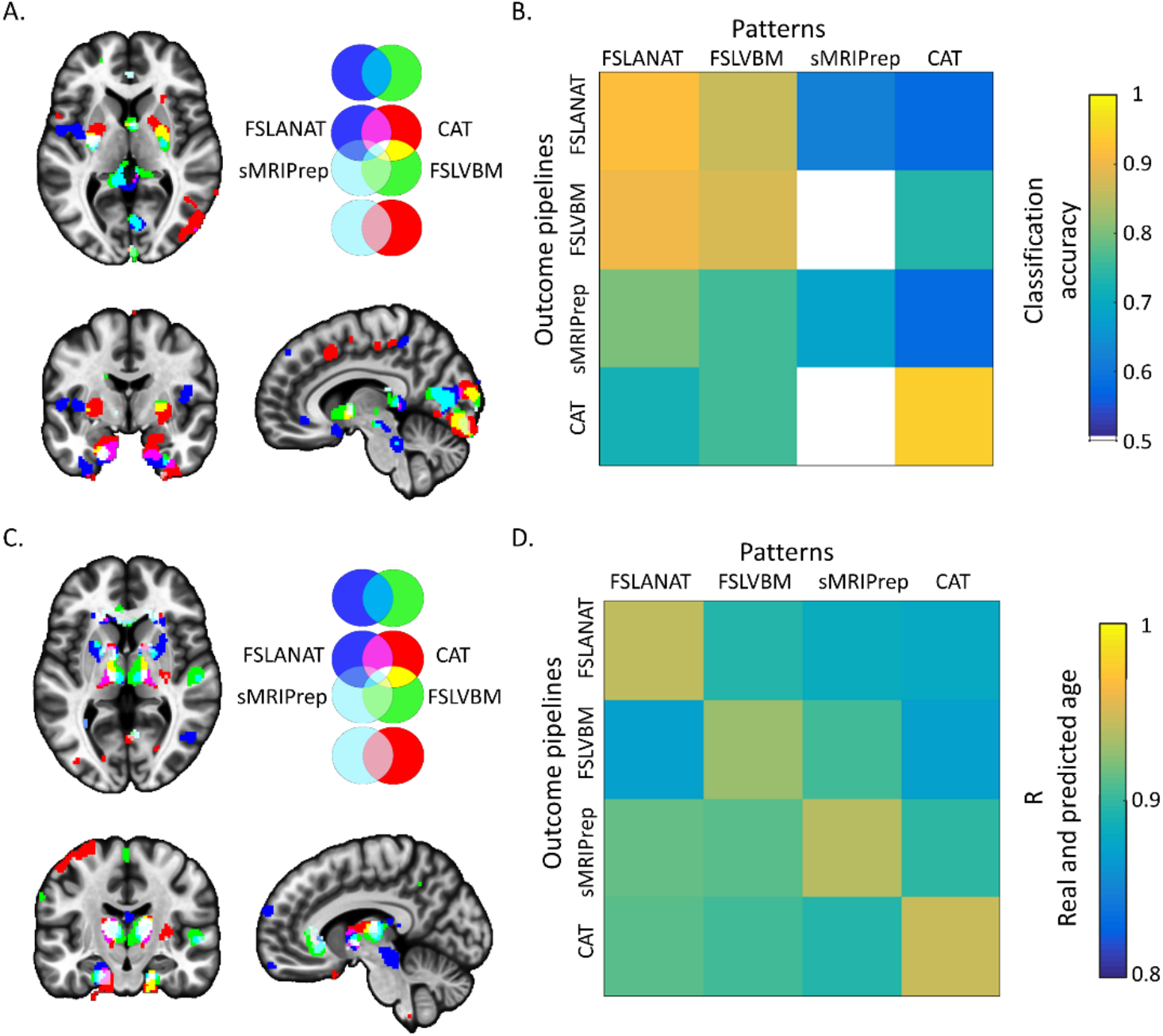
Reliable brain patterns and prediction performances for sex and age. (A) Reliable brain patterns to distinguish sex differences via bootstrapping test (5,000 permutations, *p_FDR_* < 0.05), and (B) crosspredicted accuracy of four pipelines on independent samples. The color from cold to warm indicates increased classification accuracy (from 0.5 to 1). (C) Reliable brain patterns to predict age via bootstrapping test (5,000 permutations, *p_FDR_* < 0.05), and (D) cross-predicted r value of four pipelines on each pipelines’ sample. The color from cold to warm indicates increasing r values (from 0.8 to 1).

The most reliable regions for the classification of sex across pipelines encompassed the medial prefrontal, subcortical, insular, occipital and parietal regions. Whereas overlapping clusters of predictive voxels across pipelines were only observed in the bilateral parahippocampal cortex (voxels of each cluster > 5, Fig. 6A) there were widespread differences in the location of predictive voxels. For instance, predictions based on CAT strongly weighted voxels in the putamen, hippocampus, middle cingulate cortex, and angular gyrus, while FSLANAT identified strongly predictive voxels in a widespread network including superior frontal cortex, orbitofrontal cortex, pre- and post-central cortex, insula, temporal pole, angular gyrus, and cerebellum. FSLVBM and sMRIPrep revealed generally similar findings to FSLANAT.

To further validate the impact of the processing pipelines on prediction accuracy in the independent dataset the classifiers from the training data of each pipeline were applied to the independent data processed by the other pipelines. Despite the low spatial overlap between the thresholded predictive maps all classifications across pipelines could accurately predict sex (58% ~ 94%, Cohen’s d = 0.1392 ~ 2.2815), with the exception of using the pattern developed on sMRIPrep to predict FSLVBM (50%, Cohen’s d = 0.2930) or CAT (14%, Cohen’s d = −1.4909) (Fig. 6B, corresponding Cohen’s d in Table S10). Specifically, cross-pipeline predictions between the FSL pipelines reached the highest accuracy (>86%), as well as relatively high accuracy for predicting data processed by sMRIPrep (FSLANAT: 80%, Cohen’s d = 0.6437, and FSLVBM: 76%, Cohen’s d = 0.7231) and CAT (FSLANAT: 72%, Cohen’s d = 0.7810, and FSLVBM: 76%, Cohen’s d = 0.7402).

The developed sex-predictive patterns were further validated on dataset 2 (Fig. S7) with an averaged classification accuracy of 66.33% (SD = 3.22, range = 61.94% ~ 71.86%). Given that the age range in dataset 2 was considerably higher than in the initial training dataset we limited the age range in dataset 2 to <=30 years, which increased classification accuracy in this sample (n = 159, female = 99) to an average of 71.82% (SD = 6.33, range = 61.64% ~ 87.42%, corresponding Cohen’s d in Table S10). The highest accuracy (87.42%, Cohen’s d = 2.2062) appeared when using the pattern from CAT on data processed by CAT pipeline, followed by the pattern developed from FSLANAT applied to data processed by sMRIPrep (77.99%, Cohen’s d = 1.2160).

### Association approach: age-related effects from univariate analyses

To additionally examine differences between the pipelines with respect to determining associations between GM and linear variables, age-related volumetric changes were computed using a regression approach. For parametric statistics with a cluster-level *p_FWE_* < 0.05 threshold, overlapping age-related increases were only observed for two pipelines (FSLVBM and sMRIPrep) with minimal overlap in the cerebellum (3.41% overlap, Fig. 5C and Table 2). Whereas all pipelines determined GMV decreases with age, overlap between all pipelines was only observed in the middle occipital gyrus (Fig. 5C and Table 2). Further inspection revealed that FSLANAT had rather low overlap with the other pipelines, whereas the other three pipelines additionally identified common age-related decreases in medial prefrontal, cingulate and some parietal and temporal regions (Fig. 5C). The variability of results was further reflected in both, the direction (FSLVBM and sMRIPrep), and extent of the age-related effect (sMRIPrep). Regarding non-parametric statistics with TFCE *p_FWE_* < 0.05, the results were very similar with parametric statistics, particularly for the brain regions that decreased with age (Fig. S6B and Table 2, details please see ***supplemental results***).

**Table 2.** Percent overlap of age associated GMV-changes between the pipelines

|  | Positive association |  | Negative association |  |
| --- | --- | --- | --- | --- |
|  | Parametric<sup>1</sup> | Non-parametric<sup>2</sup> | Parametric | Non-parametric |
| CAT (unique) | - | 0.52% | 21.58% | 14.35% |
| FSLVBM (unique) | 92.6% | 96.37% | 1.05% | 1.2% |
| FSLANAT (unique) | - | - | - | - |
| sMRIPrep (unique) | 3.98% | 1.87% | 4.01% | 4.25% |
| CAT ∩ FSLVBM | - | - | 0.69% | 0.77% |
| CAT ∩ FSLANAT | - | - | - | - |
| CAT ∩ sMRIPrep | - | - | 53.77% | 56.62% |
| FSLVBM ∩ FSLANAT | - | - | - | - |
| FSLVBM ∩ sMRIPrep | 3.41% | 1.23% | 0.9% | 1.05% |
| FSLANAT ∩ sMRIPrep | - | - | 0.003% | 0.003% |
| CAT ∩ FSLVBM ∩ FSLANAT | - | - | - | - |
| CAT ∩ FSLVBM ∩ sMRIPrep | - | - | 17.95% | 21.67% |
| CAT ∩ FSLANAT ∩ sMRIPrep | - | - | 0.05% | 0.07% |
| FSLVBM ∩ FSLANAT ∩ sMRIPrep | - | - | - | 0.0001% |
| CAT ∩ FSLVBM ∩ FSLANAT ∩ sMRIPrep | - | - | 0.002% | 0.02% |

<sup>1</sup>cluster-level $p_{FWE}<0.05$
<sup>2</sup>TFCE $p_{FWE}<0.05$

Further examining the spatial similarity of the age-related GMV association maps by means of correlation between unthresholded statistical maps across four pipelines revealed variations in age-related effects ranging from −0.0051 to 0.6757 (Fig. 5D). The lowest similarity values reflected that FSLANAT produced very different maps for age-related changes as compared to the other pipelines and that CAT was rather different from the FSLVBM processing pipelines. CAT and sMRIPrep had the highest similarity with respect to the unthresholded maps. In line with overlap results (Table 2), a considerable proportion of the variance between pipelines was introduced by the results from FSLANAT.

### Prediction approach: Age-related effects from multivariate pattern analysis

The overlap of pattern expressions from the four pipelines was mainly located on bilateral pallidum, bilateral thalamus, parahippocampal cortex (voxels of each cluster >5, Fig. 6C). The predicted age from four pattern expressions exhibited a high correlation with the true age of the participants (all r values >0.8, Fig. 6D). With the exception of the common subcortical regions, FSLANAT additionally revealed high predictive weight for regions in the putamen, hippocampus, hypothalamus, brainstem, medial frontal cortex, middle temporal gyrus, middle frontal gyrus and insula. In addition, FSLVBM revealed postcentral gyrus, superior frontal gyrus, superior temporal gyrus, and cerebellum, whereas CAT highlighted contributions of GMV variations in the precentral gyrus, and precuneus for age predictions.

## Discussion

The VBM approach is the most widely employed one for examining regional differences or variations in brain structure in fundamental neuroscience and psychiatric neuroimaging. We here examined the reproducibility and reliability of VBM across the most commonly used software packages and pipelines. Additionally, we examined how the choice of the processing pipeline will determine the results in terms of the identified brain regions in two prototypical VBM study scenarios examining GMV between-group differences (sex differences) or linear associations (age-related changes). To this end data from two independent datasets were processed with the recommended default options in the most widely used VBM analysis packages (CAT12, FSL, and sMRIPrep) or pipelines (FSLANAT, FSLVBM), respectively. Examining spatial similarity between the preprocessed data revealed considerable differences in the voxel-level spatial distribution of GMV across the pipelines as well as with respect to the spatial homogeneity of participants data within the pipelines. Both, voxel-level and image-based reliability analyses revealed consistently poor reliability between pipelines, confirming considerable variations in the estimation of regional GMV. We next examined how the different processing pipelines would impact the determination of GMV variations in two typical mass-univariate analytic scenarios examining between group differences (sex differences) and associations with phenotypic variations (age associations). Across parametric and non-parametric correction procedures while some overlap in the identified regions was found, there were considerable variations in both GMV sex-differences and age-associations, reflecting that the choice of softwares has a strong impact of the identified regions. In addition to mass-univariate methods, machine learning based approaches were applied to explore general associations between subtle spatial variations in GMV and the two biological variables of sex and age. Although the regional overlap of the most predictive voxels between the pipelines was low, GMV maps processed with each pipeline generally allowed an accurate prediction of the biological variables. Prediction accuracy varied within and between pipelines suggesting that the choice of processing software influences multivariate prediction accuracy. Together the findings indicate considerable variability in the results obtained and that the choice of processing pipeline will considerably influence which regions are identified in VBM analyses. This in turn will strongly influence the interpretation of the findings in terms of e.g. ‘which brain regions differ between men and women’ or ‘which brain regions show age-related declines in volume’. On the other hand, the high predictive accuracy for sex and age indicates that all GMV maps encoded biologically significant variations, although region-specific interpretations need to be considered with caution.

### Variability and reliability of the preprocessed data between the pipelines

In a first step we examined the spatial variability and reliability of the preprocessed GMV images between the pipelines. For a biological valid and robust index which reflects regional variations in gray matter one would expect a high spatial homogeneity as well as reliability across pipelines. However, the spatial similarity analyses revealed considerable variations between the pipelines as well as within them. Variability widely existed both within- and between-pipelines, but notably the samples preprocessed with CAT exhibited a higher within-participants homogeneity compared to other pipelines (Fig. 3). These findings reflect that the choice of the pipeline has a considerable influence on spatial distribution of GMV variations and additionally influences how much individual variation between individuals is retained after preprocessing of the data. Examination of the between-pipeline reliability revealed a generally poor consistency reflecting a low inter-pipeline reliability, with the additional voxel-level reliability examination suggesting some regional variations with particularly low consistency between the pipelines in parietal and frontal cortices (Fig. 4, Table S11).

### Effects of pipeline on the results of univariate analyses

Our second main aim was to examine how the choice of pipeline and implementation of the pipeline-specific default configuration would affect the results of a typical VBM study. A recent study has revealed that structural brain-behavior associations appear not to be robust for latent psychological variables while associations with biological variables such as age exhibit a higher robustness (Fjell et al., 2014; Kharabian Masouleh et al., 2016; Kharabian Masouleh et al., 2019; Lotze et al., 2019; Marek et al., 2020; Ritchie et al., 2018; Willette & Kapogiannis, 2015). With respect to GMV variations and biological factors sex and age have been extensively examined in previous studies. Although the specific regions that exhibit GMV differences between men and women differ between the studies, the regional specific differences are commonly interpreted to underlie sex-differences in cognitive and emotional functions associated with the specific regions identified (Lotze et al., 2019; Ritchie et al., 2018; Ruigrok et al., 2014). Similarly, previous findings on regional-specific GMV changes with age were divergent (Fjell et al., 2014; Kharabian Masouleh et al., 2019) - even with strongly increasing sample sizes (Chen et al., 2007; Good et al., 2001; Lotze et al., 2019; Ritchie et al., 2018) - which are commonly interpreted to reflect atrophic changes that mediate specific emotional and cognitive changes with age. In contrast, the present findings indicate that the specific regions that are identified in both, sex-differences as well as age-related changes, strongly depend on the choice of the processing pipeline. For instance, after controlling for the influence of statistical inference (same statistical software, see also similarity of unthresholded maps, Fig. 5) and sample or scanner differences (same dataset was used across pipelines) only few – or in the case of sex differences even no (Fig. 5A, Table 1) – overlapping regions were identified. Moreover, for identified GMV sex-differences no three pipelines overlapped more than 20 percent which also reflected the high regional variations. For instance, after processing with CAT results would indicate higher GM volume in men in limbic regions which are typically associated with emotional processes or spatial navigation, whereas results for the FSL-based pipeline would indicate GMV sex-differences in posterior parietal regions typically associated with attention or motor integration. With respect to age-related changes the pipelines revealed some overlapping regional GMV decreases in the middle occipital gyrus although this was generally small (0.002% and 0.02% corresponding to parametric and non-parametric statistics respectively, Fig. 5C, Fig. S6, Table 2). In general, the location, extent and direction of age-related GMV changes differed considerably between the pipelines. For instance, while the analysis with CAT revealed widespread age-related GMV decline in nearly the entire cortex, FSLANAT revealed rather regional-specific decreases in the inferior frontal regions and in contrast FSLVBM revealed regional-specific age-related GMV increases in the cerebellar and limbic regions. In line with a recent study examining the influence of pipelines on functional brain activation results (Botvinik-Nezer et al., 2020), we additionally examined spatial correlations between the unthresholded statistical maps. However, although this previous study reported a considerable overlap of the unthresholded functional maps (Botvinik-Nezer et al., 2020) cross-pipeline overlap for the GMV maps in the present study was rather low (Fig. 5), implying that the impact of pipeline additionally varies by the brain modality under investigation.

### Consistency and inconsistency in machine learning based multivariate analyses across pipelines

In addition to mass-univariate analyses, machine learning based approaches were employed to investigate sex differences and age-related effects from a functional and general biological validity perspective (Hebart & Baker, 2018; Kohoutova et al., 2020). Briefly, the basic idea is that some features that can be derived from the GMV maps significantly contribute to the accurate prediction of the biological variables age and sex. Notably, based on all GMV maps reliable features for an accurate prediction of the biological variables could be extracted (e.g. for age all correlations >0.8, Fig. 6D, for sex classifiers, higher than chance level, Fig. 6B and Fig. S7). These results suggest that from a functional perspective all pipelines retained biologically and functionally relevant information. However, further examination of the spatial distribution of the most predictive voxels revealed considerable variations across the four pipelines similar to the mass-univariate analyses (Fig. 6A and Fig. 6C). For instance, the application of CAT processed data to develop sex-classifiers would have emphasized the regional-specific contribution of the putamen, hippocampus, middle cingulate cortex, and angular gyrus, while FSLANAT would have emphasized that a widespread distributed pattern allowed successful sex classification. Finally, the preprocessing pipeline had a significant effect on prediction accuracy and prediction effect sizes, such that depending on the pipeline our sex classifiers reached 70-94% classification accuracy in an independent dataset, which indicates that the processing pipeline can have a considerable effect on the sensitivity and specificity of multivariate predictive signatures.

### The challenge of interpretability

Our findings challenge the reproducibility as well as biologically and functional interpretability of regional GMV variations as determined by the VBM approach. The choice of software had considerable impact on the regional variation of GMV on the voxel level which is difficult to reconcile with a biologically valid index. Moreover, regions that were found to exhibit sex-differences or age-related GMV changes differed strongly depending on the pipeline employed. The high variability in the regions identified would have led to a rather different functional interpretation of sex-differences (e.g. Fig. 5E) as well as atrophic changes with age and potential functional consequences. In contrast, multivariate analyses accurately predicted age and gender with classifiers trained on GMV maps from all pipelines, however, the specific predictive regions differed. Together, these findings indicate that gray matter volume indices encode biologically relevant information, yet the interpretation of specific regions in both univariate as well as multivariate analyses, cannot be drawn. In the context of the replicability crisis, meta-analyses of neuroimaging data are considered as the gold-standard, but our current findings indicate that coordinate-based meta-analyses need to account for regional variability between studies introduced by the use of different pipelines.

### Future directions

The present findings emphasize the need for detailed reporting of the software specifications and configurations which is also advocated by the Committee on Best Practices in Data Analysis and Sharing (COBIDAS) report (Nichols et al., 2017). However, the fact that the pipelines with recommended default configurations revealed completely different GMV results argues for further efforts to address the reliability after reproducibility crisis (Botvinik-Nezer et al., 2020; Hong et al., 2019; Milkowski et al., 2018). Potential initial steps are open cooperation and replicability analyses across software platforms and pipelines, open software platforms that allow comparisons and standardization of methods across platforms and a transparent and detailed processing report that should accompany manuscript submissions (e.g. as provided by sMRIPrep and fMRIPrep). Finally, the neuroimaging-based brain volumetric procedures need to be benchmarked with clear biological indices from animal models, postmortem analysis or invasive approaches.

## Conclusion

The present study demonstrated considerable variations in GMV indices and corresponding results across the most commonly used processing pipelines for VBM. The combination of mass-univariate analyses and machine learning based multivariate approaches revealed that the specific regions identified to exhibit GMV sex-differences or age-related changes varied strongly depending on the software chosen. While multivariate prediction of sex and age was possible across pipelines, prediction accuracy varied strongly between them. Together the findings challenge the interpretability and robustness of VBM results.

## References

Ashburner, J., & Friston, K. J. (2000). Voxel-based morphometry--them ethods. Neuroimage, 11(6 Pt 1), 805–821. https://doi.org/10.1006/nimg.2000.0582

Avants, B. B., Epstein, C. L., Grossman, M., & Gee, J. C. (2008). Symmetric diffeomorphic image registration with cross-correlation: evaluating automated labeling of elderly and neurodegenerative brain. Med Image Anal, 12(1), 26–41. https://doi.org/10.1016/j.media.2007.06.004

Becker, B., Wagner, D., Koester, P., Tittgemeyer, M., Mercer-Chalmers-Bender, K., Hurlemann, R., Zhang, J., Gouzoulis-Mayfrank, E., Kendrick, K. M., & Daumann, J. (2015). Smaller amygdala and medial prefrontal cortex predict escalating stimulant use. Brain, 138(Pt 7), 2074–2086. https://doi.org/10.1093/brain/awv113

Bookstein, F. L. (2001). “Voxel-based morphometry” should not be used with imperfectly registered images. Neuroimage, 14(6), 1454–1462. https://doi.org/10.1006/nimg.2001.0770

Botvinik-Nezer, R., Holzmeister, F., Camerer, C. F., Dreber, A., Huber, J., Johannesson, M., Kirchler, M., Iwanir, R., Mumford, J. A., Adcock, R. A., Avesani, P., Baczkowski, B. M., Bajracharya, A., Bakst, L., Ball, S., Barilari, M., Bault, N., Beaton, D., Beitner, J., Benoit, R. G., Berkers, R., Bhanji, J. P., Biswal, B. B., Bobadilla-Suarez, S., Bortolini, T., Bottenhorn, K. L., Bowring, A., Braem, S., Brooks, H. R., Brudner, E. G., Calderon, C. B., Camilleri, J. A., Castrellon, J. J., Cecchetti, L., Cieslik, E. C., Cole, Z. J., Collignon, O., Cox, R.W., Cunningham, W. A., Czoschke, S., Dadi, K., Davis, C. P., Luca, A., Delgado, M. R., Demetriou, L., Dennison, J. B., Di, X., Dickie, E. W., Dobryakova, E., Donnat, C. L., Dukart, J., Duncan, N. W., Durnez, J., Eed, A., Eickhoff, S. B., Erhart, A., Fontanesi, L., Fricke, G. M., Fu, S., Galvan, A., Gau, R., Genon, S., Glatard, T., Glerean, E., Goeman, J. J., Golowin, S. A. E., Gonzalez-Garcia, C., Gorgolewski, K. J., Grady, C. L., Green, M. A., Guassi Moreira, J. F., Guest, O., Hakimi, S., Hamilton, J. P., Hancock, R., Handjaras, G., Harry, B. B., Hawco, C., Herholz, P., Herman, G., Heunis, S., Hoffstaedter, F., Hogeveen, J., Holmes, S., Hu, C. P., Huettel, S. A., Hughes, M. E., Iacovella, V., Iordan, A. D., Isager, P. M., Isik, A. I., Jahn, A., Johnson, M. R., Johnstone, T., Joseph, M. J. E., Juliano, A. C., Kable, J. W., Kassinopoulos, M., Koba, C., Kong, X. Z., Koscik, T. R., Kucukboyaci, N. E., Kuhl, B. A., Kupek, S., Laird, A. R., Lamm, C., Langner, R., Lauharatanahirun, N., Lee, H., Lee, S., Leemans, A., Leo, A., Lesage, E., Li, F., Li, M. Y. C., Lim, P. C., Lintz, E. N., Liphardt, S.W., Losecaat Vermeer, A. B., Love, B. C., Mack, M. L., Malpica, N., Marins, T., Maumet, C., McDonald, K., McGuire, J. T., Melero, H., Mendez Leal, A. S., Meyer, B., Meyer, K. N., Mihai, G., Mitsis, G. D., Moll, J., Nielson, D. M., Nilsonne, G., Notter, M. P., Olivetti, E., Onicas, A. I., Papale, P., Patil, K. R., Peelle, J. E., Perez, A., Pischedda, D., Poline, J. B., Prystauka, Y., Ray, S., Reuter-Lorenz, P. A., Reynolds, R. C., Ricciardi, E., Rieck, J. R., Rodriguez-Thompson, A. M., Romyn, A., Salo, T., Samanez-Larkin, G. R., Sanz-Morales, E., Schlichting, M. L., Schultz, D. H., Shen, Q., Sheridan, M. A., Silvers, J. A., Skagerlund, K., Smith, A., Smith, D. V., Sokol-Hessner, P., Steinkamp, S. R., Tashjian, S. M., Thirion, B., Thorp, J. N., Tinghog, G., Tisdall, L., Tompson, S. H., Toro-Serey, C., Torre Tresols, J. J., Tozzi, L., Truong, V., Turella, L., van ‘t Veer, A. E., Verguts, T., Vettel, J. M., Vijayarajah, S., Vo, K., Wall, M. B., Weeda, W. D., Weis, S., White, D. J., Wisniewski, D., Xifra-Porxas, A., Yearling, E. A., Yoon, S., Yuan, R., Yuen, K. S. L., Zhang, L., Zhang, X., Zosky, J. E., Nichols, T. E., Poldrack, R. A., & Schonberg, T. (2020). Variability in the analysis of a single neuroimaging dataset by many teams. Nature, 582(7810), 84–88. https://doi.org/10.1038/s41586-020-2314-9

Buimer, E. E. L., Pas, P., Brouwer, R. M., Froeling, M., Hoogduin, H., Leemans, A., Luijten, P., van Nierop, B. J., Raemaekers, M., Schnack, H. G., Teeuw, J., Vink, M., Visser, F., Hulshoff Pol, H. E., & Mandl, R. C. W. (2020). The YOUth cohort study: MRI protocol and test-retest reliability in adults. Dev Cogn Neurosci, 45, 100816. https://doi.org/10.1016/j.dcn.2020.100816

Chen, X., Sachdev, P. S., Wen, W., & Anstey, K. J. (2007). Sex differences in regional gray matter in healthy individuals aged 44-48 years: a voxel-based morphometric study. Neuroimage, 36(3), 691–699. https://doi.org/10.1016/j.neuroimage.2007.03.063

Cicchetti, D. V., & Sparrow, S. A. (1981). Developing criteria for establishing interrater reliability of specific items: applications to assessment of adaptive behavior. Am J Ment Defic, 86(2), 127–137. https://www.ncbi.nlm.nih.gov/pubmed/7315877

Esteban, O., Birman, D., Schaer, M., Koyejo, O. O., Poldrack, R. A., & Gorgolewski, K. J. (2017). MRIQC: Advancing the automatic prediction of image quality in MRI from unseen sites. PLoS One, 12(9), e0184661. https://doi.org/10.1371/journal.pone.0184661

Esteban, O., Markiewicz, C. J., Blair, R. W., Moodie, C. A., Isik, A. I., Erramuzpe, A., Kent, J. D., Goncalves, M., DuPre, E., Snyder, M., Oya, H., Ghosh, S. S., Wright, J., Durnez, J., Poldrack, R. A., & Gorgolewski, K. J. (2019). fMRIPrep: a robust preprocessing pipeline for functional MRI. Nat Methods, 16(1), 111–116. https://doi.org/10.1038/s41592-018-0235-4

Fjell, A. M., Westlye, L. T., Grydeland, H., Amlien, I., Espeseth, T., Reinvang, I., Raz, N., Dale, A. M., Walhovd, K. B., & Alzheimer Disease Neuroimaging, I. (2014). Accelerating cortical thinning: unique to dementia or universal in aging? Cereb Cortex, 24(4), 919–934. https://doi.org/10.1093/cercor/bhs379

Gennatas, E. D., Avants, B. B., Wolf, D. H., Satterthwaite, T. D., Ruparel, K., Ciric, R., Hakonarson, H., Gur, R. E., & Gur, R. C. (2017). Age-Related Effects and Sex Differences in Gray Matter Density, Volume, Mass, and Cortical Thickness from Childhood to Young Adulthood. J Neurosci, 37(20), 5065–5073. https://doi.org/10.1523/JNEUROSCI.3550-16.2017

Good, C. D., Johnsrude, I. S., Ashburner, J., Henson, R. N., Friston, K. J., & Frackowiak, R. S. (2001). A voxel-based morphometric study of ageing in 465 normal adult human brains. Neuroimage, 14(1 Pt 1), 21–36. https://doi.org/10.1006/nimg.2001.0786

Gorgolewski, K., Burns, C. D., Madison, C., Clark, D., Halchenko, Y. O., Waskom, M. L., & Ghosh, S. S. (2011). Nipype: a flexible, lightweight and extensible neuroimaging data processing framework in python. Front Neuroinform, 5, 13. https://doi.org/10.3389/fninf.2011.00013

Hebart, M. N., & Baker, C. I. (2018). Deconstructing multivariate decoding for the study of brain function. Neuroimage, 180(Pt A), 4–18. https://doi.org/10.1016/j.neuroimage.2017.08.005

Hong, Y. W., Yoo, Y., Han, J., Wager, T. D., & Woo, C. W. (2019). False-positive neuroimaging: Undisclosed flexibility in testing spatial hypotheses allows presenting anything as a replicated finding. Neuroimage, 195, 384–395. https://doi.org/10.1016/j.neuroimage.2019.03.070

Jenkinson, M., Beckmann, C. F., Behrens, T. E., Woolrich, M. W., & Smith, S. M. (2012). Fsl. Neuroimage, 62(2), 782–790. https://doi.org/10.1016/j.neuroimage.2011.09.015

Kharabian Masouleh, S., Arelin, K., Horstmann, A., Lampe, L., Kipping, J. A., Luck, T., Riedel-Heller, S. G., Schroeter, M. L., Stumvoll, M., Villringer, A., & Witte, A. V. (2016). Higher body mass index in older adults is associated with lower gray matter volume: implications for memory performance. Neurobiol Aging, 40, 1–10. https://doi.org/10.1016/j.neurobiolaging.2015.12.020

Kharabian Masouleh, S., Eickhoff, S. B., Hoffstaedter, F., Genon, S., & Alzheimer’s Disease Neuroimaging, I. (2019). Empirical examination of the replicability of associations between brain structure and psychological variables. Elife, 8. https://doi.org/10.7554/eLife.43464

Kharabian Masouleh, S., Eickhoff, S. B., Zeighami, Y., Lewis, L. B., Dahnke, R., Gaser, C., Chouinard-Decorte, F., Lepage, C., Scholtens, L. H., Hoffstaedter, F., Glahn, D. C., Blangero, J., Evans, A. C., Genon, S., & Valk, S. L. (2020). Influence of Processing Pipeline on Cortical Thickness Measurement. Cereb Cortex, 30(9), 5014–5027. https://doi.org/10.1093/cercor/bhaa097

Kohoutova, L., Heo, J., Cha, S., Lee, S., Moon, T., Wager, T. D., & Woo, C. W. (2020). Toward a unified framework for interpreting machine-learning models in neuroimaging. Nat Protoc, 15(4), 1399–1435. https://doi.org/10.1038/s41596-019-0289-5

Lakens, D. (2013). Calculating and reporting effect sizes to facilitate cumulative science: a practical primer for t-tests and ANOVAs. Front Psychol, 4, 863. https://doi.org/10.3389/fpsyg.2013.00863

Liu, C., Xu, L., Li, J., Zhou, F., Yang, X., Zheng, X., Fu, M., Li, K., Sindermann, C., Montag, C., Ma, Y., Scheele, D., Ebstein, R. P., Yao, S., Kendrick, K. M., & Becker, B. (2020). Serotonin and early life stress interact to shape brain architecture and anxious avoidant behavior - a TPH2 imaging genetics approach. Psychol Med, 1–9. https://doi.org/10.1017/S0033291720002809

Llera, A., Wolfers, T., Mulders, P., & Beckmann, C. F. (2019). Inter-individual differences in human brain structure and morphology link to variation in demographics and behavior. Elife, 8. https://doi.org/10.7554/eLife.44443

Lotze, M., Domin, M., Gerlach, F. H., Gaser, C., Lueders, E., Schmidt, C. O., & Neumann, N. (2019). Novel findings from 2,838 Adult Brains on Sex Differences in Gray Matter Brain Volume. Sci Rep, 9(1), 1671. https://doi.org/10.1038/s41598-018-38239-2

Madan, C. R., & Kensinger, E. A. (2017). Test-retest reliability of brain morphology estimates. Brain Inform, 4(2), 107–121. https://doi.org/10.1007/s40708-016-0060-4

Marek, S., Tervo-Clemmens, B., Calabro, F. J., Montez, D. F., Kay, B. P., Hatoum, A. S., Donohue, M. R., Foran, W., Miller, R. L., Feczko, E., Miranda-Dominguez, O., Graham, A. M., Earl, E. A., Perrone, A. J., Cordova, M., Doyle, O., Moore, L. A., Conan, G., Uriarte, J., Snider, K., Tam, A., Chen, J., Newbold, D. J., Zheng, A., Seider, N. A., Van, A. N., Laumann, T. O., Thompson, W. K., Greene, D. J., Petersen, S. E., Nichols, T. E., Yeo, B. T. T., Barch, D. M., Garavan, H., Luna, B., Fair, D. A., & Dosenbach, N. U. F. (2020). Towards Reproducible Brain-Wide Association Studies. https://doi.org/10.1101/2020.08.21.257758

Mechelli, A., Friston, K. J., Frackowiak, R. S., & Price, C. J. (2005). Structural covariance in the human cortex. J Neurosci, 25(36), 8303–8310. https://doi.org/10.1523/JNEUROSCI.0357-05.2005

Melzer, T. R., Keenan, R. J., Leeper, G. J., Kingston-Smith, S., Felton, S. A., Green, S. K., Henderson, K. J., Palmer, N. J., Shoorangiz, R., Almuqbel, M. M., & Myall, D. J. (2020). Test-retest reliability and sample size estimates after MRI scanner relocation. Neuroimage, 211, 116608. https://doi.org/10.1016/j.neuroimage.2020.116608

Milkowski, M., Hensel, W. M., & Hohol, M. (2018). Replicability or reproducibility? On the replication crisis in computational neuroscience and sharing only relevant detail. J Comput Neurosci, 45(3), 163–172. https://doi.org/10.1007/s10827-018-0702-z

Miller, K. L., Alfaro-Almagro, F., Bangerter, N. K., Thomas, D. L., Yacoub, E., Xu, J., Bartsch, A. J., Jbabdi, S., Sotiropoulos, S. N., Andersson, J. L., Griffanti, L., Douaud, G., Okell, T. W., Weale, P., Dragonu, I., Garratt, S., Hudson, S., Collins, R., Jenkinson, M., Matthews, P. M., & Smith, S. M. (2016). Multimodal population brain imaging in the UK Biobank prospective epidemiological study. Nat Neurosci, 19(11), 1523–1536. https://doi.org/10.1038/nn.4393

Nichols, T. E., Das, S., Eickhoff, S. B., Evans, A. C., Glatard, T., Hanke, M., Kriegeskorte, N., Milham, M. P., Poldrack, R. A., Poline, J. B., Proal, E., Thirion, B., Van Essen, D. C., White, T., & Yeo, B. T. (2017). Best practices in data analysis and sharing in neuroimaging using MRI. Nat Neurosci, 20(3), 299–303. https://doi.org/10.1038/nn.4500

Noble, S., Scheinost, D., & Constable, R. T. (2019). A decade of test-retest reliability of functional connectivity: A systematic review and meta-analysis. Neuroimage, 203, 116157. https://doi.org/10.1016/j.neuroimage.2019.116157

Noble, S., Spann, M. N., Tokoglu, F., Shen, X., Constable, R. T., & Scheinost, D. (2017). Influences on the Test-Retest Reliability of Functional Connectivity MRI and its Relationship with Behavioral Utility. Cereb Cortex, 27(11), 5415–5429. https://doi.org/10.1093/cercor/bhx230

Nostro, A. D., Muller, V. I., Reid, A. T., & Eickhoff, S. B. (2017). Correlations Between Personality and Brain Structure: A Crucial Role of Gender. Cereb Cortex, 27(7), 3698–3712. https://doi.org/10.1093/cercor/bhw191

Nunes, A., Schnack, H. G., Ching, C. R. K., Agartz, I., Akudjedu, T. N., Alda, M., Alnaes, D., Alonso-Lana, S., Bauer, J., Baune, B. T., Boen, E., Bonnin, C. D. M., Busatto, G. F., Canales-Rodriguez, E. J., Cannon, D. M., Caseras, X., Chaim-Avancini, T. M., Dannlowski, U., Diaz-Zuluaga, A. M., Dietsche, B., Doan, N. T., Duchesnay, E., Elvsashagen, T., Emden, D., Eyler, L. T., Fatjo-Vilas, M., Favre, P., Foley, S. F., Fullerton, J. M., Glahn, D. C., Goikolea, J. M., Grotegerd, D., Hahn, T., Henry, C., Hibar, D. P., Houenou, J., Howells, F. M., Jahanshad, N., Kaufmann, T., Kenney, J., Kircher, T. T. J., Krug, A., Lagerberg, T. V., Lenroot, R. K., Lopez-Jaramillo, C., Machado-Vieira, R., Malt, U. F., McDonald, C., Mitchell, P. B., Mwangi, B., Nabulsi, L., Opel, N., Overs, B. J., Pineda-Zapata, J. A., Pomarol-Clotet, E., Redlich, R., Roberts, G., Rosa, P. G., Salvador, R., Satterthwaite, T. D., Soares, J. C., Stein, D. J., Temmingh, H. S., Trappenberg, T., Uhlmann, A., van Haren, N. E. M., Vieta, E., Westlye, L. T., Wolf, D. H., Yuksel, D., Zanetti, M. V., Andreassen, O. A., Thompson, P. M., Hajek, T., & Group, E. B. D. W. (2020). Using structural MRI to identify bipolar disorders - 13 site machine learning study in 3020 individuals from the ENIGMA Bipolar Disorders Working Group. Mol Psychiatry, 25(9), 2130–2143. https://doi.org/10.1038/s41380-018-0228-9

Pepe, A., Dinov, I., & Tohka, J. (2014). An automatic framework for quantitative validation of voxel based morphometry measures of anatomical brain asymmetry. Neuroimage, 100, 444–459. https://doi.org/10.1016/j.neuroimage.2014.06.029

Poldrack, R. A., Baker, C. I., Durnez, J., Gorgolewski, K. J., Matthews, P. M., Munafo, M. R., Nichols, T. E., Poline, J. B., Vul, E., & Yarkoni, T. (2017). Scanning the horizon: towards transparent and reproducible neuroimaging research. Nat Rev Neurosci, 18(2), 115–126. https://doi.org/10.1038/nrn.2016.167

Popescu, V., Schoonheim, M. M., Versteeg, A., Chaturvedi, N., Jonker, M., Xavier de Menezes, R., Gallindo Garre, F., Uitdehaag, B. M., Barkhof, F., & Vrenken, H. (2016). Grey Matter Atrophy in Multiple Sclerosis: Clinical Interpretation Depends on Choice of Analysis Method. PLoS One, 11(1), e0143942. https://doi.org/10.1371/journal.pone.0143942

Premi, E., Cauda, F., Costa, T., Diano, M., Gazzina, S., Gualeni, V., Alberici, A., Archetti, S., Magoni, M., Gasparotti, R., Padovani, A., & Borroni, B. (2016). Looking for Neuroimaging Markers in Frontotemporal Lobar Degeneration Clinical Trials: A Multi-Voxel Pattern Analysis Study in Granulin Disease. J Alzheimers Dis, 51(1), 249–262. https://doi.org/10.3233/JAD-150340

Rajagopalan, V., & Pioro, E. P. (2015). Disparate voxel based morphometry (VBM) results between SPM and FSL softwares in ALS patients with frontotemporal dementia: which VBM results to consider? BMC Neurol, 15, 32. https://doi.org/10.1186/s12883-015-0274-8

Ritchie, S. J., Cox, S. R., Shen, X., Lombardo, M. V., Reus, L. M., Alloza, C., Harris, M. A., Alderson, H. L., Hunter, S., Neilson, E., Liewald, D. C. M., Auyeung, B., Whalley, H. C., Lawrie, S. M., Gale, C. R., Bastin, M. E., McIntosh, A. M., & Deary, I. J. (2018). Sex Differences in the Adult Human Brain: Evidence from 5216 UK Biobank Participants. Cereb Cortex, 28(8), 2959–2975. https://doi.org/10.1093/cercor/bhy109

Ruigrok, A. N., Salimi-Khorshidi, G., Lai, M. C., Baron-Cohen, S., Lombardo, M. V., Tait, R. J., & Suckling, J. (2014). A meta-analysis of sex differences in human brain structure. Neurosci Biobehav Rev, 39, 34–50. https://doi.org/10.1016/j.neubiorev.2013.12.004

Senjem, M. L., Gunter, J. L., Shiung, M. M., Petersen, R. C., & Jack, C. R., Jr. (2005). Comparison of different methodological implementations of voxel-based morphometry in neurodegenerative disease. Neuroimage, 26(2), 600–608. https://doi.org/10.1016/j.neuroimage.2005.02.005

Shen, S., Szameitat, A. J., & Sterr, A. (2007). VBM lesion detection depends on the normalization template: a study using simulated atrophy. Magnetic Resonance Imaging, 25(10), 1385–1396. https://doi.org/10.1016/j.mri.2007.03.025

Shou, H., Eloyan, A., Lee, S., Zipunnikov, V., Crainiceanu, A. N., Nebel, N. B., Caffo, B., Lindquist, M. A., & Crainiceanu, C. M. (2013). Quantifying the reliability of image replication studies: the image intraclass correlation coefficient (I2C2). Cogn Affect Behav Neurosci, 13(4), 714–724. https://doi.org/10.3758/s13415-013-0196-0

Smith, S. M., Jenkinson, M., Woolrich, M. W., Beckmann, C. F., Behrens, T. E., Johansen-Berg, H., Bannister, P. R., De Luca, M., Drobnjak, I., Flitney, D. E., Niazy, R. K., Saunders, J., Vickers, J., Zhang, Y., De Stefano, N., Brady, J. M., & Matthews, P. M. (2004). Advances in functional and structural MR image analysis and implementation as FSL. Neuroimage, 23 Suppl 1, S208–219. https://doi.org/10.1016/j.neuroimage.2004.07.051

Taubert, M., Lohmann, G., Margulies, D. S., Villringer, A., & Ragert, P. (2011). Long-term effects of motor training on resting-state networks and underlying brain structure. Neuroimage, 57(4), 1492–1498. https://doi.org/10.1016/j.neuroimage.2011.05.078

Tustison, N. J., Avants, B. B., Cook, P. A., Zheng, Y., Egan, A., Yushkevich, P. A., & Gee, J. C. (2010). N4ITK: improved N3 bias correction. IEEE Trans Med Imaging, 29(6), 1310–1320. https://doi.org/10.1109/TMI.2010.2046908

Wang, Q., Chen, C., Cai, Y., Li, S., Zhao, X., Zheng, L., Zhang, H., Liu, J., Chen, C., & Xue, G. (2016). Dissociated neural substrates underlying impulsive choice and impulsive action. Neuroimage, 134, 540–549. https://doi.org/10.1016/j.neuroimage.2016.04.010

Wei, D., Zhuang, K., Ai, L., Chen, Q., Yang, W., Liu, W., Wang, K., Sun, J., & Qiu, J. (2018). Structural and functional brain scans from the cross-sectional Southwest University adult lifespan dataset. Sci Data, 5, 180134. https://doi.org/10.1038/sdata.2018.134

Willette, A. A., & Kapogiannis, D. (2015). Does the brain shrink as the waist expands? Ageing Res Rev, 20, 86–97. https://doi.org/10.1016/j.arr.2014.03.007

Yamashita, A., Yahata, N., Itahashi, T., Lisi, G., Yamada, T., Ichikawa, N., Takamura, M., Yoshihara, Y., Kunimatsu, A., Okada, N., Yamagata, H., Matsuo, K., Hashimoto, R., Okada, G., Sakai, Y., Morimoto, J., Narumoto, J., Shimada, Y., Kasai, K., Kato, N., Takahashi, H., Okamoto, Y., Tanaka, S. C., Kawato, M., Yamashita, O., & Imamizu, H. (2019). Harmonization of resting-state functional MRI data across multiple imaging sites via the separation of site differences into sampling bias and measurement bias. PLoS Biol, 17(4), e3000042. https://doi.org/10.1371/journal.pbio.3000042

Yan, C. G., Wang, X. D., Zuo, X. N., & Zang, Y. F. (2016). DPABI: Data Processing & Analysis for (Resting-State) Brain Imaging. Neuroinformatics, 14(3), 339–351. https://doi.org/10.1007/s12021-016-9299-4

Zhang, Y., Brady, M., & Smith, S. (2001). Segmentation of brain MR images through a hidden Markov random field model and the expectation-maximization algorithm. IEEE Trans Med Imaging, 20(1), 45–57. https://doi.org/10.1109/42.906424

Zhao, Z., Yao, S., Zweerings, J., Zhou, X., Zhou, F., Kendrick, K. M., Chen, H., Mathiak, K., & Becker, B. (2021). Putamen volume predicts real-time fMRI neurofeedback learning success across paradigms and neurofeedback target regions. Hum Brain Mapp. https://doi.org/10.1002/hbm.25336

Zhou, X., Wu, R., Liu, C., Kou, J., Chen, Y., Pontes, H. M., Yao, D., Kendrick, K. M., Becker, B., & Montag, C. (2020). Higher levels of (Internet) Gaming Disorder symptoms according to the WHO and APA frameworks associate with lower striatal volume. J Behav Addict, 9(3), 598–605. https://doi.org/10.1556/2006.2020.00066

