## supplemental material for "Location, location, location– choice of Voxel-Based Morphometry processing pipeline drives variability in the location of neuroanatomical brain markers"

Zhou et al.

**Supplemental Figures S1-S8**
**Supplemental Tables S1-S11**

**Correspondence:**
Benjamin Becker,
Xinqi Zhou,
Center for Information in Medicine
University of Electronic Science and Technology of China
Chengdu 611731, China

### Supplemental methods

*Structural MRI acquisition parameters*

Both datasets were acquired using validated T1-weighted brain structural acquisition protocols. Dataset 1 was acquired on a 3.0 Tesla GE MR750 system (General Electric Medical Systems, Milwaukee, WI, USA). T1-weighted high-resolution anatomical images were acquired with a spoiled gradient echo pulse sequence, repetition time (TR) = 5.9 ms, echo time (TE) = 2 ms, flip angle = 9°, field of view (FOV) = 256 × 256 mm, acquisition matrix = 256 × 256, thickness = 1 mm, number of slices = 156, voxel size=1×1×1 mm. Dataset 2 was collected using a 3.0-T Siemens Trio MRI scanner (Siemens Medical, Erlangen, Germany). A magnetization-prepared rapid gradient echo (MPRAGE) sequence was used to acquire high-resolution T1-weighted anatomical images (repetition time = 1,900 ms, echo time = 2.52 ms, inversion time = 900 ms, flip angle = 90°, resolution matrix = 256 × 256, slices = 176, thickness = 1.0 mm, and voxel size = 1×1×1 mm) (details see also Wei et al., 2018).

*Overlapping percent of mass-univariate analyses*

To estimate the consistency in terms of spatial overlap between the pipelines percent overlapping voxels were calculated for the pipeline-specific results on sex differences and age-related changes respectively (Table S10 & S11). The following formula was applied:

$$p_i = \frac{v_i}{v_{all}} \times 100$$

$p_i$ indicates the overlapping percent of $i^{th}$ overlapping or independent cluster voxels ($v_i$) in voxels of all significant results ($v_{all}$), which represents the union set of the four pipelines.

### Supplemental results

*Spatial similarity within- and between-pipelines*

In the male sample, the ANOVA revealed that main effects of pipeline were significant with respect to all pipeline and sample homogeneity comparisons, including between pipelines and between participants (F = 4635, $p$ < 0.0001), between pipelines and within participants (F = 179, $p$ < 0.0001), and within pipelines and between participants (F = 14208, $p$ < 0.0001). The post hoc tests were conducted with appropriate Bonferroni correction (Table S1, S3, and S5).

In the female sample, the ANOVA revealed that main effects of pipeline were significant with respect to all pipeline and sample homogeneity comparisons, including between pipelines and between participants (F = 3535, $p$ <0.0001), between pipelines and within participants (F = 196.1, $p$ <0.0001), and within pipelines and between participants (F = 9894, $p$ <0.0001). The post hoc tests were conducted with appropriate Bonferroni correction (Table S2, S4, and S6).

For dataset 2, the ANOVA revealed that main effects of pipeline for all homogeneity comparisons, including between pipelines and between participants (F = 61376, $p$ <0.0001), between pipelines and within participants (F = 593.7, $p$ <0.0001), and within pipelines and between participants (F = 290745, $p$ <0.0001). Post hoc tests were conducted with Bonferroni's correction (Table S7, S8, and S9).

*Between-group approach: sex differences univariate analyses*

Results from the non-parametric statistics with TFCE $p_{FWE} < 0.05$ were highly similar to the parametric statistic results, suggesting that the pipeline differences are robust across statistic models. For instance, across pipelines males had higher GMV than females (FSLANAT, FSLVBM, and CAT had 18.89% overlaps, Table 1) in the precuneus, bilateral putamen, insula, olfactory cortex, parahippocampal cortex, and cerebellum (Fig.S6A, while for the FSL pipelines (FSLANAT and FSLVBM had 9.33% overlap, Table 1) females had higher GMV in inferior parietal lobule, postcentral cortex, and angular gyrus. Again, the software packages revealed widespread differences with respect to sex-differences in GMV in limbic, frontal and cerebellar regions. Notably, in some instances the overlap between the software packages increased slightly using the non-parametric approach (Table 1 and Fig. S6A).

*Association approach: age-related effects from univariate analyses*

Regarding to non-parametric statistics with TFCE $p_{FWE} < 0.05$, the results were very similar with parametric statistics, especially for the brain regions that decreased with age. FSLVBM revealed age-related increases from prefrontal cortex to parietal lobe to cerebellum, and bilateral hippocampus, while sMRIPrep revealed caudate and cerebellum. In addition, CAT highlighted thalamus, but only FSLVBM and sMRIPrep identified common cerebellar regions that increased with age (Fig. S6B and Table 2). Whereas CAT and sMRIPrep revealed age-related decreases in widespread regions covering nearly the entire cortex, the other pipelines revealed more regional-specific decreases with age, such that FSLANAT revealed specific decreases in the inferior frontal gyrus and middle occipital gyrus. FSLVBM additionally revealed regional decreases in middle cingulate cortex, frontal and temporal cortex. Again, the common brain regions across four pipelines that decreased with age only included the middle occipital gyrus (Fig. S6B). Except for FSLANAT the common regions of the other pipelines included medial prefrontal cortex, cingulate gyrus, precuneus, temporal lobe, parietal lobe, middle occipital gyrus, insula, and cerebellum (Fig. S6B).

### Supplemental figures

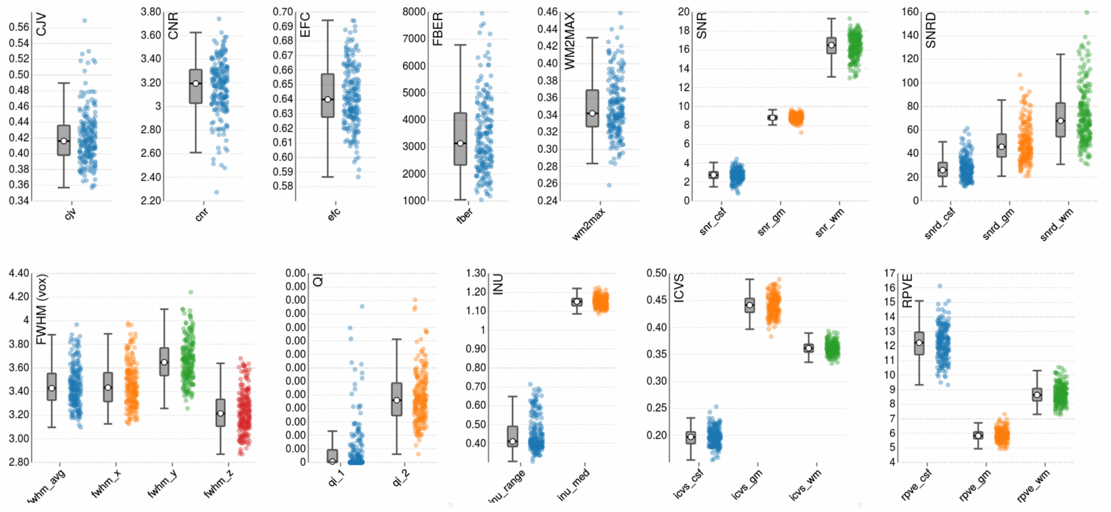


Fig. S1. Quality metrics for dataset 1.

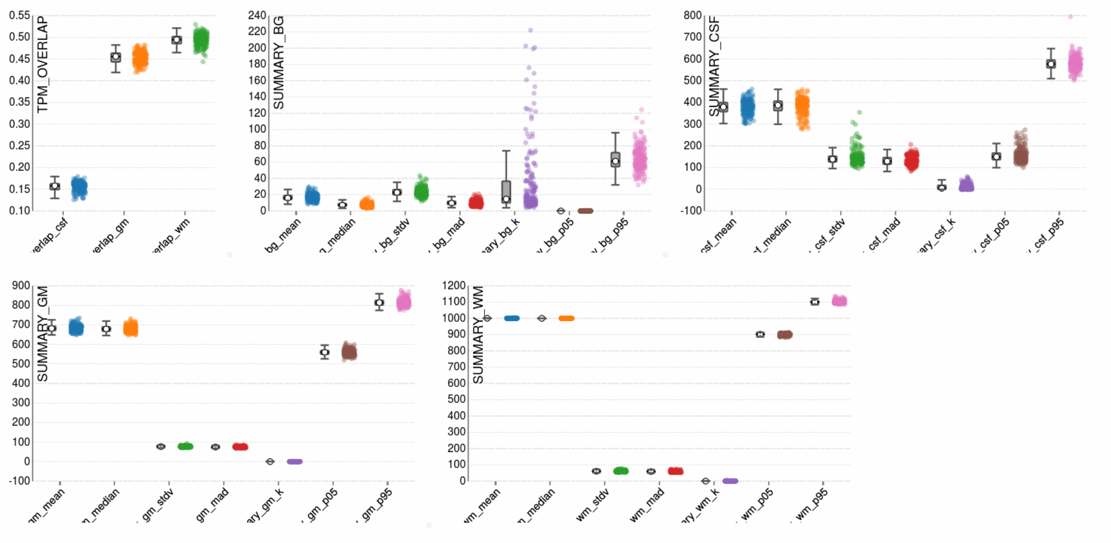


Fig. S2. Quality metrics for dataset 1.

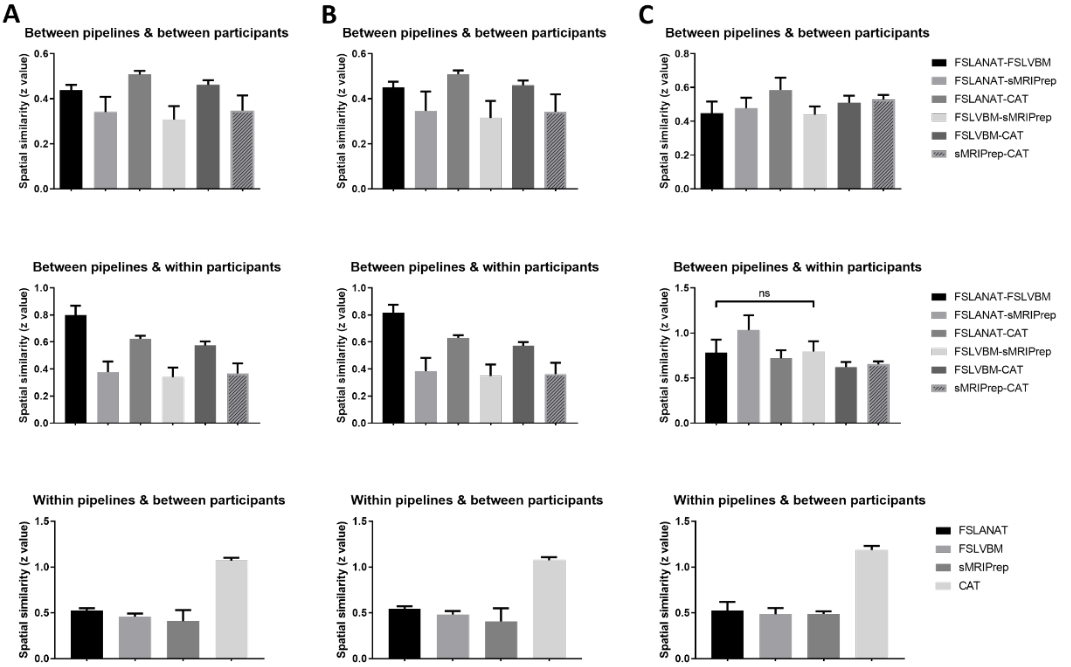


Fig. S3. The mean similarity and SD of different pipelines and pipelines' pairs for the unsmoothed data. The column A represents males, column B represents females, from dataset 1; column C represents dataset 2.

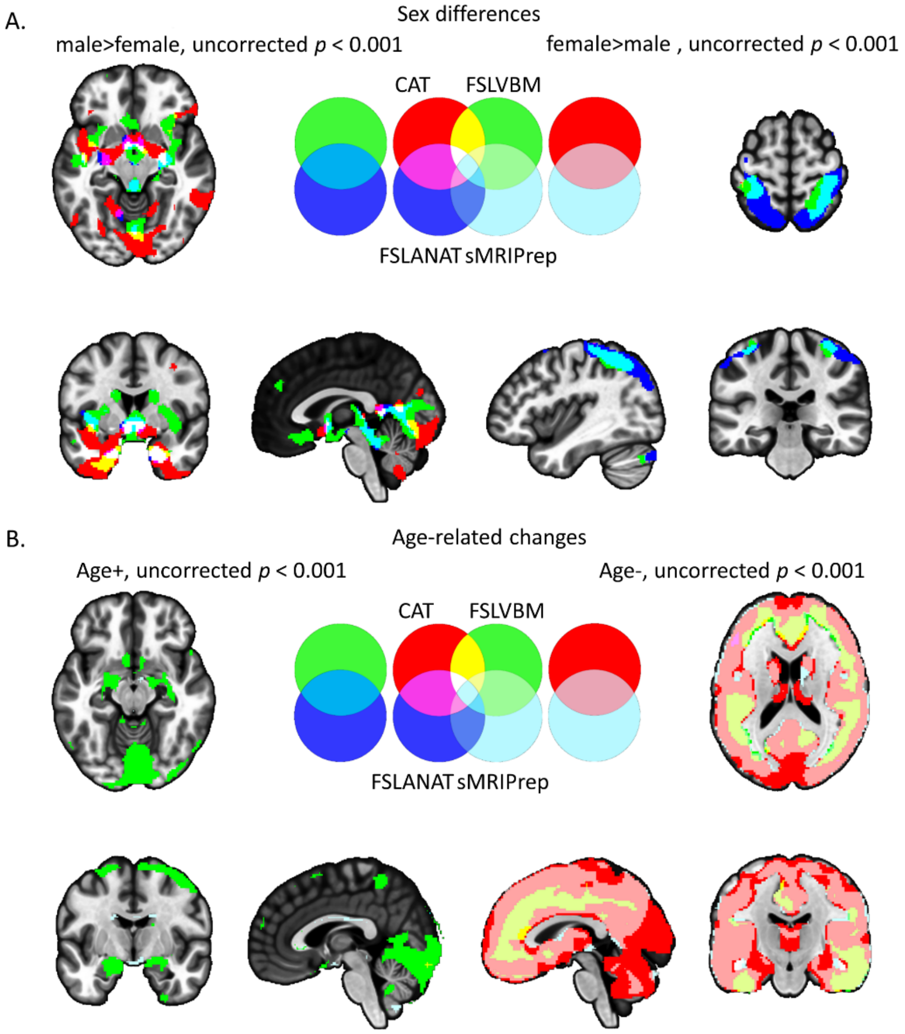


Fig. S4. Similarities and dissimilarities between the pipelines with respect to determining GMV sex differences and age-related GMV changes. Displayed in A and B are results from parametric statistic overlaps at uncorrected $p < 0.001$. The left panels of A display results for the male>female contrast. The right panels of A correspond to the female>male contrast. The left panels of B depicts brain regions with increasing GMV with age. The right panels of B depict decreases with age. Red = CAT, green = FSLVBM, blue = FSLANAT, light blue = sMRIPrep, other colors visualize the overlap between the results.

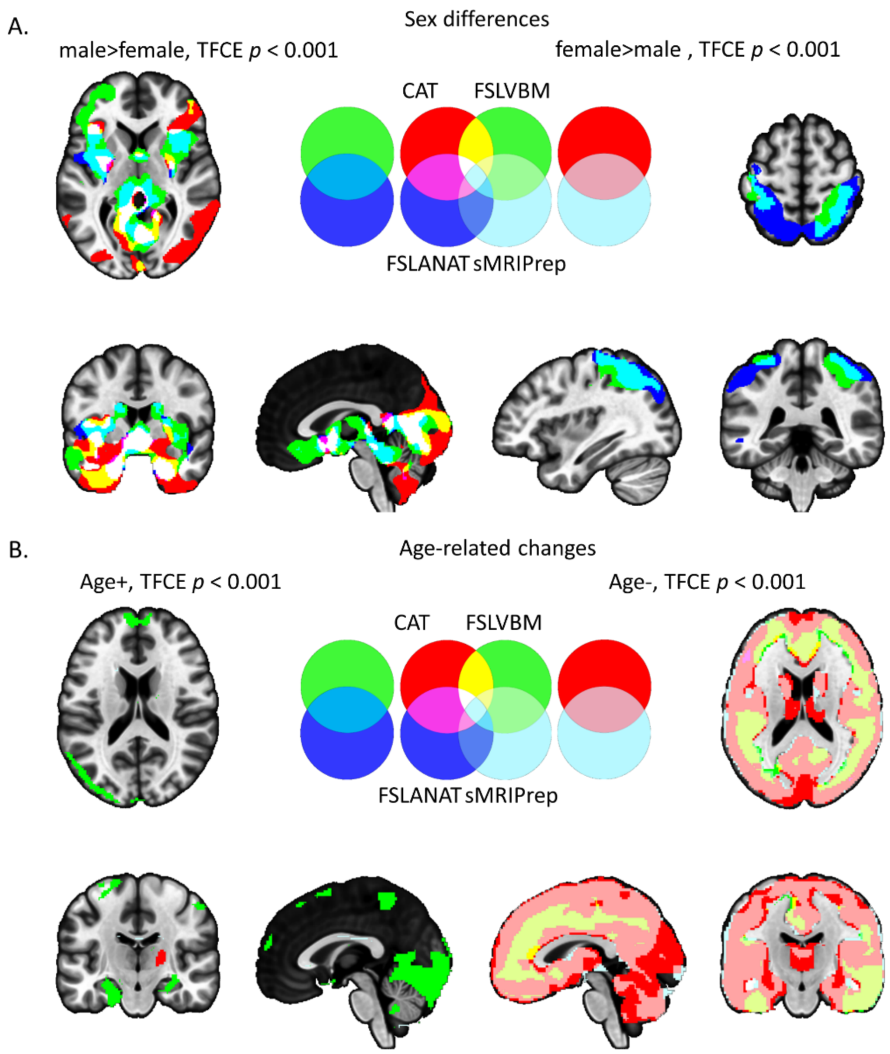


Fig. S5. Similarities and dissimilarities between the pipelines with respect to determining GMV sex differences. Displayed of A and B are results from non-parametric statistics (TFCE with 5,000 permutations) overlapping at $p < 0.001$. The left panels of A display results for the male>female contrast. The right panels of A correspond to the female>male contrast. The left panels of B depicts brain regions with increasing GMV with age. The right panels of B depict decreases with age. Red = CAT, green = FSLVBM, blue = FSLANAT, light blue = sMRIPrep, other colors visualize the overlap between the results.

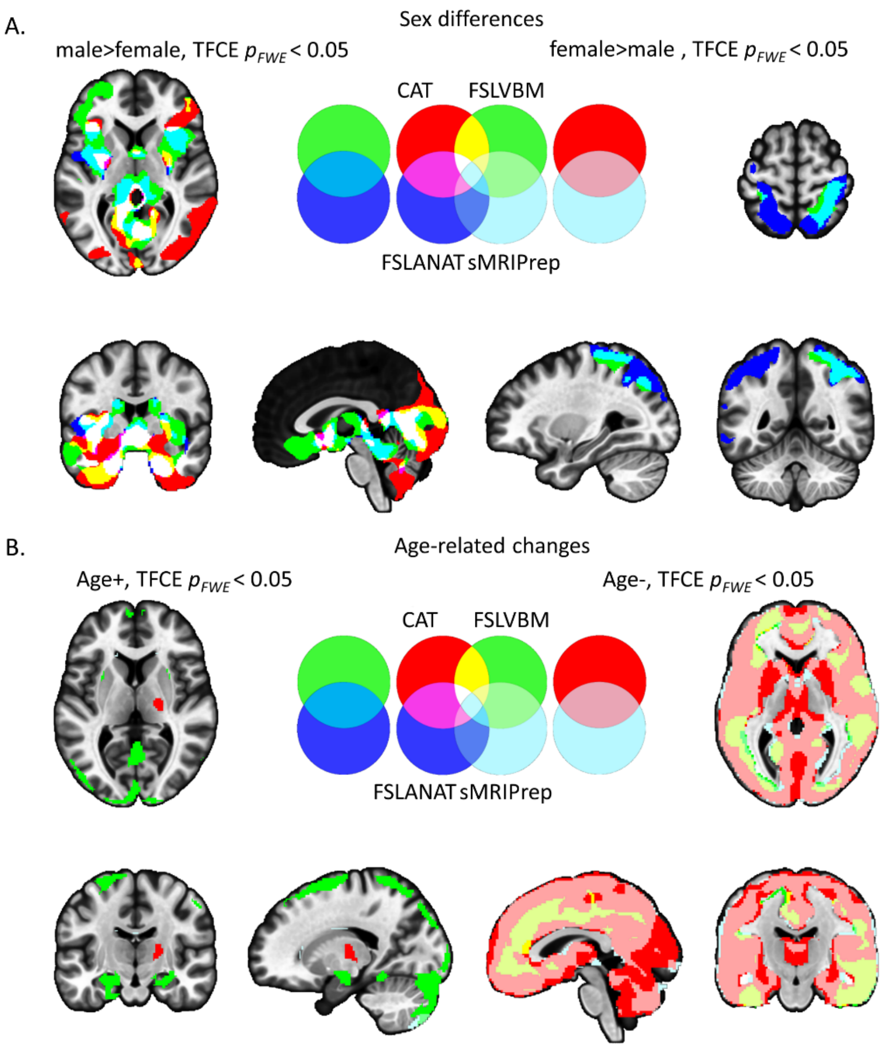


Fig. S6. Similarities and dissimilarities between the pipelines with respect to determining GMV sex differences. Displayed of A and B are results from non-parametric statistics (TFCE with 5,000 permutations) overlapping at $p_{FWE}$ < 0.05. The left panels of A display results for the male>female contrast. The right panels of A correspond to the female>male contrast. The left panels of B depicts brain regions with increasing GMV with age. The right panels of B depict decreases with age. Red = CAT, green = FSLVBM, blue = FSLANAT, light blue = sMRIPrep, other colors visualize the overlap between the results.

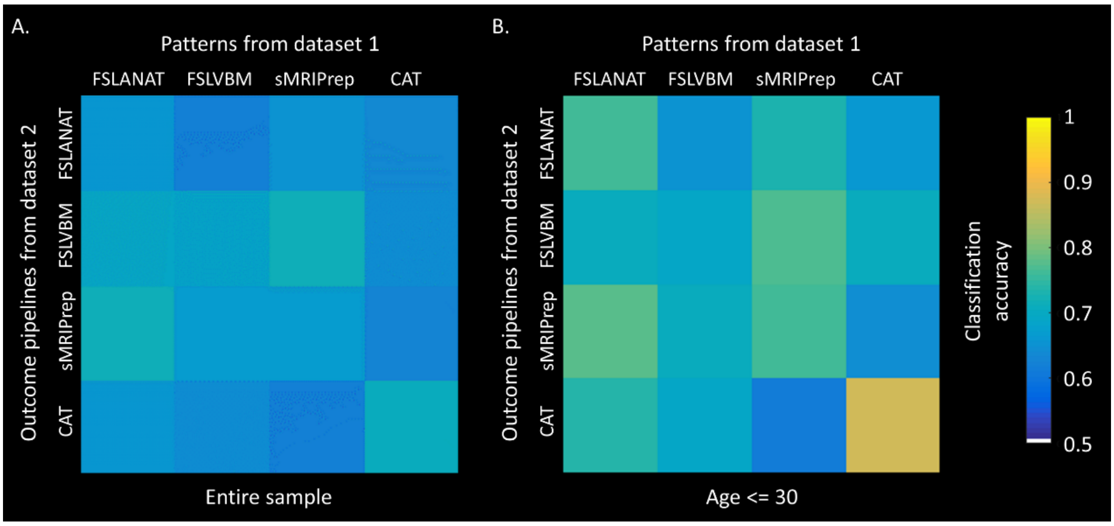


Fig. S7. Classification accuracy of patterns from dataset 1 on dataset 2 for (A) the entire sample, and (B) only age <= 30 (n = 159, female = 99).

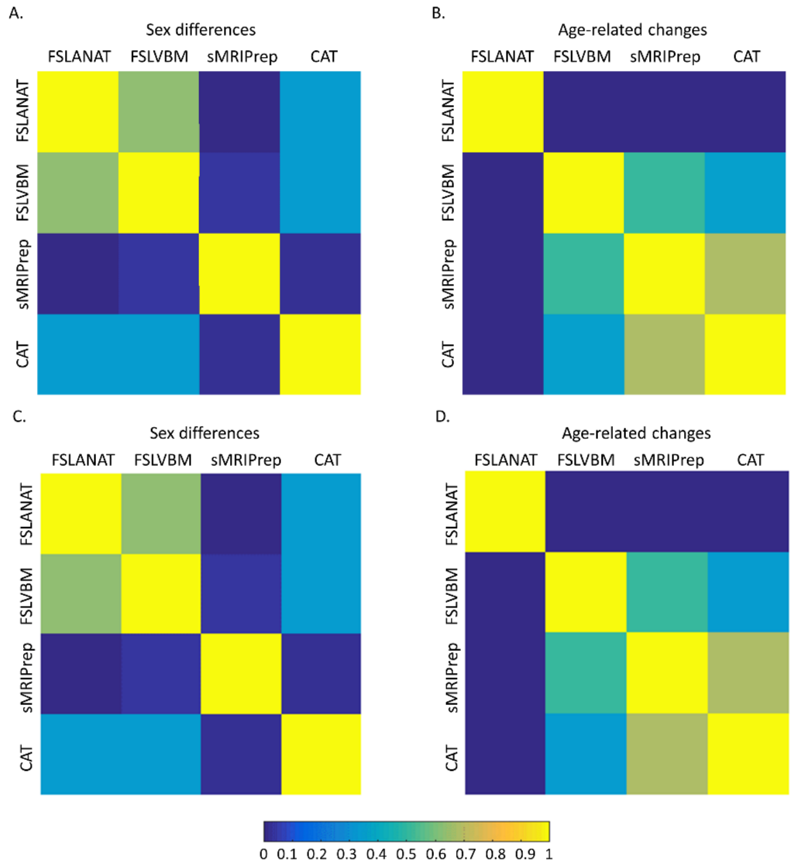


Fig. S8. Variability of unthresholded statistical maps. The correlation values between whole-brain unthresholded statistical maps of four pipelines were computed respectively for (A, C) sex differences, and (B, D) age-related effects. The different MNI templates do not affect the results between A/B using East Asian template, and C/D using Caucasian template. Only positive values are showed for display purpose.

**Supplemental tables**

Table S1. Multiple comparisons for between-pipelines and between-participants spatial similarity of male

| Bonferroni's multiple comparisons test | Mean Diff. | 95.00% CI of diff. | Adjusted P Value |
|---|---|---|---|
| FSLANAT-FSLVBM vs. FSLANAT-sMRIPrep | 0.09092 | 0.08494 to 0.09690 | <0.0001 |
| FSLANAT-FSLVBM vs. FSLANAT-CAT | -0.03750 | -0.04013 to -0.03487 | <0.0001 |
| FSLANAT-FSLVBM vs. FSLVBM-sMRIPrep | 0.1520 | 0.1461 to 0.1578 | <0.0001 |
| FSLANAT-FSLVBM vs. FSLVBM-CAT | 0.03990 | 0.03669 to 0.04311 | <0.0001 |
| FSLANAT-FSLVBM vs. sMRIPrep-CAT | 0.1809 | 0.1752 to 0.1866 | <0.0001 |
| FSLANAT-sMRIPrep vs. FSLANAT-CAT | -0.1284 | -0.1339 to -0.1229 | <0.0001 |
| FSLANAT-sMRIPrep vs. FSLVBM-sMRIPrep | 0.06105 | 0.05878 to 0.06332 | <0.0001 |
| FSLANAT-sMRIPrep vs. FSLVBM-CAT | -0.05102 | -0.05684 to -0.04520 | <0.0001 |
| FSLANAT-sMRIPrep vs. sMRIPrep-CAT | 0.08998 | 0.08254 to 0.09741 | <0.0001 |
| FSLANAT-CAT vs. FSLVBM-sMRIPrep | 0.1895 | 0.1841 to 0.1948 | <0.0001 |
| FSLANAT-CAT vs. FSLVBM-CAT | 0.07740 | 0.07531 to 0.07949 | <0.0001 |
| FSLANAT-CAT vs. sMRIPrep-CAT | 0.2184 | 0.2133 to 0.2235 | <0.0001 |
| FSLVBM-sMRIPrep vs. FSLVBM-CAT | -0.1121 | -0.1171 to -0.1070 | <0.0001 |
| FSLVBM-sMRIPrep vs. sMRIPrep-CAT | 0.02893 | 0.02166 to 0.03620 | <0.0001 |
| FSLVBM-CAT vs. sMRIPrep-CAT | 0.1410 | 0.1356 to 0.1464 | <0.0001 |

Table S2. Multiple comparisons for between-pipelines and between-participants spatial similarity of female

| Bonferroni's multiple comparisons test | Mean Diff. | 95.00% CI of diff. | Adjusted P Value |
|---|---|---|---|
| FSLANAT-FSLVBM vs. FSLANAT-sMRIPrep | 0.1005 | 0.09391 to 0.1071 | <0.0001 |
| FSLANAT-FSLVBM vs. FSLANAT-CAT | 0.007626 | 0.005233 to 0.01002 | <0.0001 |
| FSLANAT-FSLVBM vs. FSLVBM-sMRIPrep | 0.1463 | 0.1401 to 0.1525 | <0.0001 |
| FSLANAT-FSLVBM vs. FSLVBM-CAT | 0.07866 | 0.07588 to 0.08144 | <0.0001 |
| FSLANAT-FSLVBM vs. sMRIPrep-CAT | 0.2164 | 0.2105 to 0.2224 | <0.0001 |
| FSLANAT-sMRIPrep vs. FSLANAT-CAT | -0.09289 | -0.09950 to -0.08628 | <0.0001 |
| FSLANAT-sMRIPrep vs. FSLVBM-sMRIPrep | 0.04578 | 0.04396 to 0.04759 | <0.0001 |
| FSLANAT-sMRIPrep vs. FSLVBM-CAT | -0.02185 | -0.02864 to -0.01506 | <0.0001 |
| FSLANAT-sMRIPrep vs. sMRIPrep-CAT | 0.1159 | 0.1074 to 0.1245 | <0.0001 |
| FSLANAT-CAT vs. FSLVBM-sMRIPrep | 0.1387 | 0.1325 to 0.1448 | <0.0001 |
| FSLANAT-CAT vs. FSLVBM-CAT | 0.07104 | 0.06956 to 0.07251 | <0.0001 |
| FSLANAT-CAT vs. sMRIPrep-CAT | 0.2088 | 0.2034 to 0.2142 | <0.0001 |
| FSLVBM-sMRIPrep vs. FSLVBM-CAT | -0.06763 | -0.07371 to -0.06155 | <0.0001 |
| FSLVBM-sMRIPrep vs. sMRIPrep-CAT | 0.07014 | 0.06203 to 0.07825 | <0.0001 |
| FSLVBM-CAT vs. sMRIPrep-CAT | 0.1378 | 0.1322 to 0.1434 | <0.0001 |

Table S3. Multiple comparisons for between-pipelines and within-participants spatial similarity of male

| Bonferroni's multiple comparisons test | Mean Diff. | 95.00% CI of diff. | Adjusted P Value |
|---|---|---|---|
| FSLANAT-FSLVBM vs. FSLANAT-sMRIPrep | 0.3494 | 0.2624 to 0.4364 | <0.0001 |
| FSLANAT-FSLVBM vs. FSLANAT-CAT | 0.1633 | 0.1119 to 0.2146 | <0.0001 |
| FSLANAT-FSLVBM vs. FSLVBM-sMRIPrep | 0.4362 | 0.3638 to 0.5087 | <0.0001 |
| FSLANAT-FSLVBM vs. FSLVBM-CAT | 0.2816 | 0.2475 to 0.3157 | <0.0001 |
| FSLANAT-FSLVBM vs. sMRIPrep-CAT | 0.5125 | 0.4334 to 0.5916 | <0.0001 |
| FSLANAT-sMRIPrep vs. FSLANAT-CAT | -0.1861 | -0.2567 to -0.1155 | <0.0001 |
| FSLANAT-sMRIPrep vs. FSLVBM-sMRIPrep | 0.08682 | 0.05555 to 0.1181 | <0.0001 |
| FSLANAT-sMRIPrep vs. FSLVBM-CAT | -0.06780 | -0.1431 to 0.007462 | 0.1190 |
| FSLANAT-sMRIPrep vs. sMRIPrep-CAT | 0.1631 | 0.1453 to 0.1809 | <0.0001 |
| FSLANAT-CAT vs. FSLVBM-sMRIPrep | 0.2729 | 0.2039 to 0.3419 | <0.0001 |
| FSLANAT-CAT vs. FSLVBM-CAT | 0.1183 | 0.08949 to 0.1471 | <0.0001 |
| FSLANAT-CAT vs. sMRIPrep-CAT | 0.3492 | 0.2906 to 0.4077 | <0.0001 |
| FSLVBM-sMRIPrep vs. FSLVBM-CAT | -0.1546 | -0.2189 to -0.09033 | <0.0001 |
| FSLVBM-sMRIPrep vs. sMRIPrep-CAT | 0.07626 | 0.04582 to 0.1067 | <0.0001 |
| FSLVBM-CAT vs. sMRIPrep-CAT | 0.2309 | 0.1671 to 0.2946 | <0.0001 |

Table S4. Multiple comparisons for between-pipelines and within-participants spatial similarity of female

| Bonferroni's multiple comparisons test | Mean Diff. | 95.00% CI of diff. | Adjusted P Value |
|---|---|---|---|
| FSLANAT-FSLVBM vs. FSLANAT-sMRIPrep | 0.3709 | 0.2862 to 0.4557 | <0.0001 |
| FSLANAT-FSLVBM vs. FSLANAT-CAT | 0.2375 | 0.2022 to 0.2728 | <0.0001 |
| FSLANAT-FSLVBM vs. FSLVBM-sMRIPrep | 0.4454 | 0.3737 to 0.5170 | <0.0001 |
| FSLANAT-FSLVBM vs. FSLVBM-CAT | 0.3520 | 0.3282 to 0.3759 | <0.0001 |
| FSLANAT-FSLVBM vs. sMRIPrep-CAT | 0.5745 | 0.5072 to 0.6418 | <0.0001 |
| FSLANAT-sMRIPrep vs. FSLANAT-CAT | -0.1335 | -0.2144 to -0.05250 | <0.0001 |
| FSLANAT-sMRIPrep vs. FSLVBM-sMRIPrep | 0.07445 | 0.05105 to 0.09784 | <0.0001 |
| FSLANAT-sMRIPrep vs. FSLVBM-CAT | -0.01890 | -0.1022 to 0.06441 | >0.9999 |
| FSLANAT-sMRIPrep vs. sMRIPrep-CAT | 0.2036 | 0.1796 to 0.2276 | <0.0001 |
| FSLANAT-CAT vs. FSLVBM-sMRIPrep | 0.2079 | 0.1324 to 0.2834 | <0.0001 |
| FSLANAT-CAT vs. FSLVBM-CAT | 0.1146 | 0.09476 to 0.1344 | <0.0001 |
| FSLANAT-CAT vs. sMRIPrep-CAT | 0.3370 | 0.2767 to 0.3974 | <0.0001 |
| FSLVBM-sMRIPrep vs. FSLVBM-CAT | -0.09335 | -0.1672 to -0.01951 | 0.0037 |
| FSLVBM-sMRIPrep vs. sMRIPrep-CAT | 0.1291 | 0.1009 to 0.1574 | <0.0001 |
| FSLVBM-CAT vs. sMRIPrep-CAT | 0.2225 | 0.1584 to 0.2865 | <0.0001 |

Table S5. Multiple comparisons for within-pipelines and between-participants spatial similarity of male

| Bonferroni's multiple comparisons test | Mean Diff. | 95.00% CI of diff. | Adjusted P Value |
|---|---|---|---|
| FSLANAT vs. FSLVBM | 0.08878 | 0.08378 to 0.09378 | <0.0001 |
| FSLANAT vs. sMRIPrep | 0.2117 | 0.2010 to 0.2223 | <0.0001 |
| FSLANAT vs. CAT | -0.4077 | -0.4105 to -0.4048 | <0.0001 |
| FSLVBM vs. sMRIPrep | 0.1229 | 0.1112 to 0.1345 | <0.0001 |
| FSLVBM vs. CAT | -0.4964 | -0.5015 to -0.4914 | <0.0001 |
| sMRIPrep vs. CAT | -0.6193 | -0.6301 to -0.6085 | <0.0001 |

Table S6. Multiple comparisons for within-pipelines and between-participants spatial similarity of female

| Bonferroni's multiple comparisons test | Mean Diff. | 95.00% CI of diff. | Adjusted P Value |
|---|---|---|---|
| FSLANAT vs. FSLVBM | 0.05479 | 0.05050 to 0.05909 | <0.0001 |
| FSLANAT vs. sMRIPrep | 0.2404 | 0.2275 to 0.2532 | <0.0001 |
| FSLANAT vs. CAT | -0.3749 | -0.3770 to -0.3727 | <0.0001 |
| FSLVBM vs. sMRIPrep | 0.1856 | 0.1720 to 0.1991 | <0.0001 |
| FSLVBM vs. CAT | -0.4297 | -0.4343 to -0.4250 | <0.0001 |
| sMRIPrep vs. CAT | -0.6152 | -0.6282 to -0.6022 | <0.0001 |

Table S7. Multiple comparisons for between-pipelines and between-participants spatial similarity of dataset 2

| Bonferroni's multiple comparisons test | Mean Diff. | 95.00% CI of diff. | Adjusted P Value |
|---|---|---|---|
| FSLANAT-FSLVBM vs. FSLANAT-sMRIPrep | -0.03518 | -0.03583 to -0.03453 | <0.0001 |
| FSLANAT-FSLVBM vs. FSLANAT-CAT | -0.1328 | -0.1336 to -0.1320 | <0.0001 |
| FSLANAT-FSLVBM vs. FSLVBM-sMRIPrep | -0.03569 | -0.03689 to -0.03449 | <0.0001 |
| FSLANAT-FSLVBM vs. FSLVBM-CAT | 0.02268 | 0.02154 to 0.02382 | <0.0001 |
| FSLANAT-FSLVBM vs. sMRIPrep-CAT | -0.01869 | -0.01971 to -0.01767 | <0.0001 |
| FSLANAT-sMRIPrep vs. FSLANAT-CAT | -0.09763 | -0.09811 to -0.09715 | <0.0001 |
| FSLANAT-sMRIPrep vs. FSLVBM-sMRIPrep | -0.0005106 | -0.001524 to 0.0005024 | >0.9999 |
| FSLANAT-sMRIPrep vs. FSLVBM-CAT | 0.05786 | 0.05688 to 0.05884 | <0.0001 |
| FSLANAT-sMRIPrep vs. sMRIPrep-CAT | 0.01649 | 0.01563 to 0.01734 | <0.0001 |
| FSLANAT-CAT vs. FSLVBM-sMRIPrep | 0.09712 | 0.09597 to 0.09827 | <0.0001 |
| FSLANAT-CAT vs. FSLVBM-CAT | 0.1555 | 0.1544 to 0.1565 | <0.0001 |
| FSLANAT-CAT vs. sMRIPrep-CAT | 0.1141 | 0.1132 to 0.1150 | <0.0001 |
| FSLVBM-sMRIPrep vs. FSLVBM-CAT | 0.05837 | 0.05797 to 0.05877 | <0.0001 |
| FSLVBM-sMRIPrep vs. sMRIPrep-CAT | 0.01700 | 0.01618 to 0.01781 | <0.0001 |
| FSLVBM-CAT vs. sMRIPrep-CAT | -0.04137 | -0.04194 to -0.04080 | <0.0001 |

Table S8. Multiple comparisons for between-pipelines and within-participants spatial similarity of dataset 2

| Bonferroni's multiple comparisons test | Mean Diff. | 95.00% CI of diff. | Adjusted P Value |
|---|---|---|---|
| FSLANAT-FSLVBM vs. FSLANAT-sMRIPrep | -0.1898 | -0.2134 to -0.1661 | <0.0001 |
| FSLANAT-FSLVBM vs. FSLANAT-CAT | 0.01048 | -0.01726 to 0.03822 | >0.9999 |
| FSLANAT-FSLVBM vs. FSLVBM-sMRIPrep | -0.1046 | -0.1304 to -0.07874 | <0.0001 |
| FSLANAT-FSLVBM vs. FSLVBM-CAT | 0.2493 | 0.2224 to 0.2762 | <0.0001 |
| FSLANAT-FSLVBM vs. sMRIPrep-CAT | 0.1713 | 0.1369 to 0.2057 | <0.0001 |
| FSLANAT-sMRIPrep vs. FSLANAT-CAT | 0.2002 | 0.1820 to 0.2185 | <0.0001 |
| FSLANAT-sMRIPrep vs. FSLVBM-sMRIPrep | 0.08520 | 0.04686 to 0.1235 | <0.0001 |
| FSLANAT-sMRIPrep vs. FSLVBM-CAT | 0.4391 | 0.4058 to 0.4724 | <0.0001 |
| FSLANAT-sMRIPrep vs. sMRIPrep-CAT | 0.3611 | 0.3304 to 0.3918 | <0.0001 |
| FSLANAT-CAT vs. FSLVBM-sMRIPrep | -0.1150 | -0.1523 to -0.07781 | <0.0001 |
| FSLANAT-CAT vs. FSLVBM-CAT | 0.2388 | 0.2098 to 0.2678 | <0.0001 |
| FSLANAT-CAT vs. sMRIPrep-CAT | 0.1608 | 0.1354 to 0.1863 | <0.0001 |
| FSLVBM-sMRIPrep vs. FSLVBM-CAT | 0.3539 | 0.3376 to 0.3702 | <0.0001 |
| FSLVBM-sMRIPrep vs. sMRIPrep-CAT | 0.2759 | 0.2479 to 0.3038 | <0.0001 |
| FSLVBM-CAT vs. sMRIPrep-CAT | -0.07800 | -0.09320 to -0.06281 | <0.0001 |

Table S9. Multiple comparisons for within-pipelines and between-participants spatial similarity of dataset 2

| Bonferroni's multiple comparisons test | Mean Diff. | 95.00% CI of diff. | Adjusted P Value |
|---|---|---|---|
| FSLANAT vs. FSLVBM | -0.05583 | -0.05793 to -0.05372 | <0.0001 |
| FSLANAT vs. sMRIPrep | -0.003483 | -0.005081 to -0.001885 | <0.0001 |
| FSLANAT vs. CAT | -0.4597 | -0.4612 to -0.4581 | <0.0001 |
| FSLVBM vs. sMRIPrep | 0.05234 | 0.05089 to 0.05379 | <0.0001 |
| FSLVBM vs. CAT | -0.4038 | -0.4053 to -0.4023 | <0.0001 |
| sMRIPrep vs. CAT | -0.4562 | -0.4567 to -0.4557 | <0.0001 |

Table S10. Cohen's d for each classification of male and female

| | FSLANAT | FSLVBM | sMRIPrep | CAT |
|---|---|---|---|---|
| Testing sample from dataset 1 | | | | |
| FSLANAT | 2.0260 | 1.4821 | 0.5963 | 0.1392 |
| FSLVBM | 1.4589 | 1.4798 | 0.2930 | 0.6472 |
| SMRIPrep | 0.6437 | 0.7231 | 0.2967 | 0.1693 |
| CAT | 0.7810 | 0.7402 | -1.4909 | 2.2815 |
| Testing sample from dataset 2 (age<=30) | | | | |
| FSLANAT | 1.0091 | 0.5780 | 0.4651 | 0.4242 |
| FSLVBM | 0.8998 | 0.8162 | 0.6787 | 0.2237 |
| SMRIPrep | 1.2160 | 0.8602 | 1.1182 | 0.2635 |
| CAT | 0.7471 | 0.7350 | -1.0130 | 2.2062 |

Table S11. Image intraclass correlation coefficient (I2C2) across pipelines

| | I2C2 | 95% CI[1] |
|---|---|---|
| Dataset 1 | | |
| FSLANAT vs. FSLVBM | 0.1802 | 0.1535 to 0.2062 |
| FSLANAT vs. sMRIPrep | 0.0489 | 0.0190 to 0.1067 |
| FSLANAT vs. CAT | 0.2983 | 0.2711 to 0.3247 |
| FSLVBM vs. sMRIPrep | 0.0695 | 0.0337 to 0.1382 |
| FSLVBM vs. CAT | 0.3750 | 0.3445 to 0.4035 |
| sMRIPrep vs. CAT | 0.0994 | 0.0574 to 0.1901 |
| Across all pipelines | 0.1613 | 0.0996 to 0.2579 |
| Dataset 2 | | |
| FSLANAT vs. FSLVBM | 0.1180 | 0.0948 to 0.1441 |
| FSLANAT vs. sMRIPrep | 0.1205 | 0.0982 to 0.1424 |
| FSLANAT vs. CAT | 0.2437 | 0.2019 to 0.2860 |
| FSLVBM vs. sMRIPrep | 0.1344 | 0.1118 to 0.1604 |
| FSLVBM vs. CAT | 0.3061 | 0.2617 to 0.3596 |
| sMRIPrep vs. CAT | 0.3439 | 0.3247 to 0.3634 |
| Across all pipelines | 0.2656 | 0.2357 to 0.2960 |

[1] 95% confidence interval implemented by 5000 bootstrapping

**References**

Wei, D., Zhuang, K., Ai, L., Chen, Q., Yang, W., Liu, W., Wang, K., Sun, J., & Qiu, J. (2018). Structural and functional brain scans from the cross-sectional Southwest University adult lifespan dataset. *Sci Data*, *5*, 180134. https://doi.org/10.1038/sdata.2018.134
